# Identification and Characterization of the First Fragment Hits for SETDB1 Tudor Domain

**DOI:** 10.1101/605303

**Authors:** Pavel Mader, Rodrigo Mendoza-Sanchez, Aman Iqbal, Aiping Dong, Elena Dobrovetsky, Victoria B. Corless, Sean K. Liew, Scott R. Houliston, Renato Ferreira De Freitas, David Smil, Carlo C. Dela Sena, Steven Kennedy, Diego B. Diaz, Hong Wu, Ludmila Dombrovski, Abdellah Allali-Hassani, Jinrong Min, Matthieu Schapira, Masoud Vedadi, Peter J. Brown, Vijayaratnam Santhakumar, Andrei K. Yudin, Cheryl H. Arrowsmith

## Abstract

SET domain bifurcated protein 1 (SETDB1) is a human histone-lysine methyltransferase, which is amplified in human cancers and was shown to be crucial in the growth of non-small and small cell lung carcinoma. In addition to its catalytic domain, SETDB1 harbors a unique tandem tudor domain which recognizes histone sequences containing both methylated and acetylated lysines, and likely contributes to its localization on chromatin. Using X-ray crystallography and NMR spectroscopy fragment screening approaches, we have identified the first small molecule fragment hits that bind to histone peptide binding groove of the TTD of SETDB1. Herein, we describe the binding modes of these fragments and analogues and the biophysical characterization of key compounds. These confirmed small molecule fragments will inform the development of potent antagonists of SETDB1 interaction with histones.

## INTRODUCTION

Post-translational modifications (PTMs) such as methylation, phosphorylation, acetylation, ubiquitination, and SUMOylation of histone proteins impact gene expression by altering chromatin compaction or recruiting effector proteins.^1-3^ Methylation of histones occurs predominantly at the lysine and arginine sidechains of histone 3 (H3K4, H3K9, H3K27, H3K36, H3K79 and H3R2) and histone 4 (H4K20 and H4R3). Unlike some other PTMs, methylation of a lysine or arginine sidechain does not change overall charge but results in subtle alterations in size and hydrophobicity, which can influence chromatin packing.^4^ Transcriptional activation or repression is dependent on the position and degree of methylation^5^ and aberrant methylation of histones can result in dysregulation of cell growth and development of cancer.^6^ Many of the histone methyl transferases have been extensively studied and selective inhibitors have been effective in preclinical models.^7-9^

Human SETDB1 is a histone-lysine methyltransferase (HMT), which specifically trimethylates histone H3 lysine 9 (H3K9me3). It is therefore deemed central in the silencing of specific euchromatic genes.^10^ Recent studies have also suggested that SETDB1 could be an oncogene. Its overexpression is crucial to the growth of non-small and small cell lung cancer cell lines and has been reported in various human cancers.^11-13^ Overexpressed SETDB1 increases H3K9me3 leading to alterations in gene expression.^14^

SETDB1 contains a tandem tudor domain (TTD), a methyl-DNA binding domain (MBD), and a SET domain.^15^ The MBD of SETDB1 promotes direct binding to methylated DNA sites and mediates trimethylation of H3K9 in a site-directed fashion. The SET domain catalytically “writes” the epigenetic methyl mark, which then goes on to recruit proteins belonging to the Heterochromatin Protein 1 (HP1) family to control gene repression.^10^ Tudor domains are associated with the recognition of different methylation levels of lysine and arginine residues and the TTD of SETDB1 has been associated with the recognition of di- and trimethylation states of H3K9.^16,17^ To the best of our knowledge, and despite the evident therapeutic importance of SETDB1, no antagonists of this protein have been reported to date. We therefore undertook fragment based approach toward discovery of ligands for SETDB1. We successfully crystallized the truncated tandem tudor domain of SETDB1 in complex with a histone 3 peptide dimethylated at lysine 9 and acetylated at lysine 14 (H3K9me2K14ac).^18^ The co-crystal structure shows that H3K9me2K14ac interacts with the protein at two interconnected pockets within the TTD: a small pocket where the methyl lysine (Kme) binds (hereafter called the Kme binding pocket) and a relatively larger and deeper pocket engaged by the acetyl lysine (the Kac pocket) (Figure 1).

**Figure 1.**
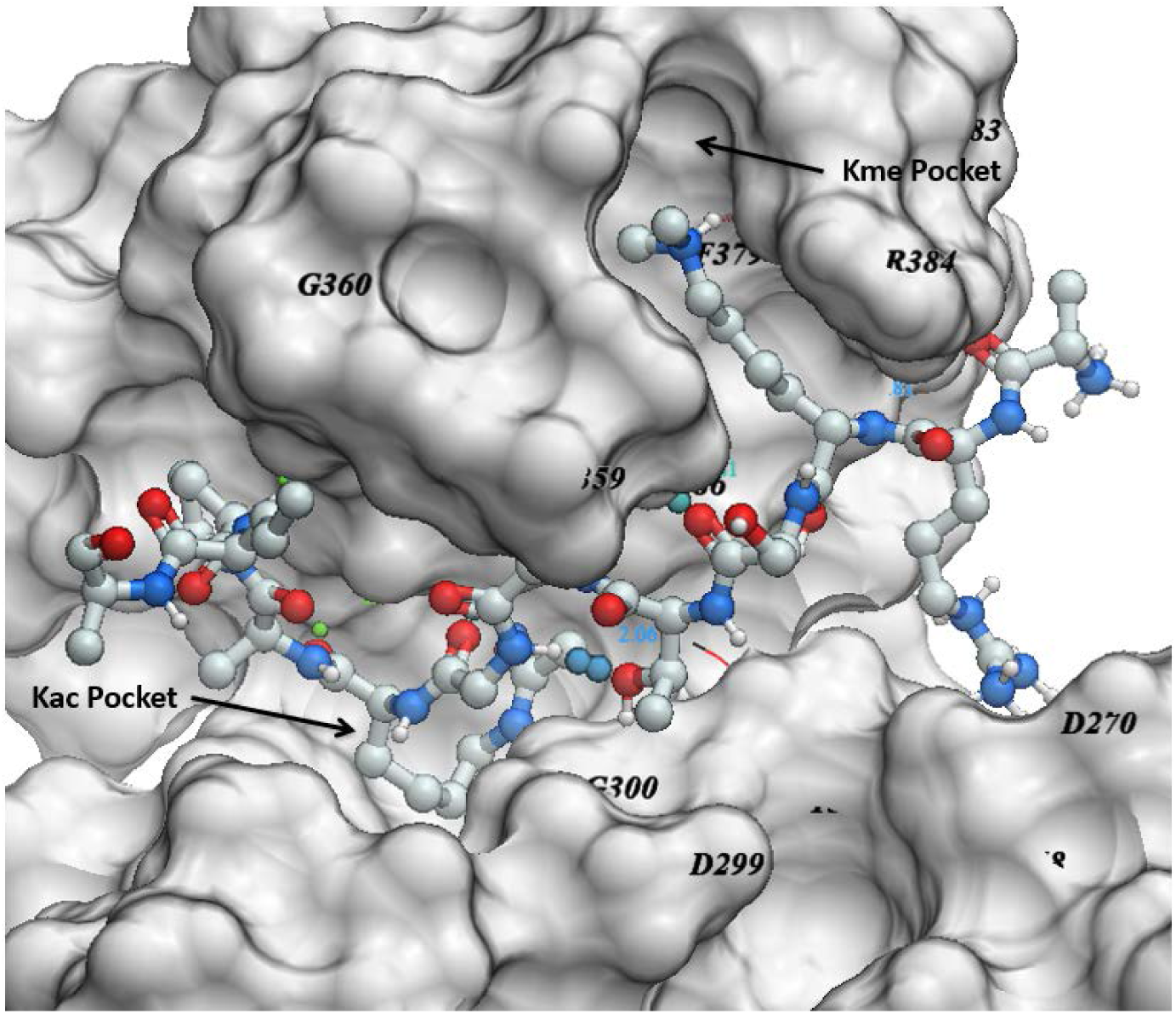
Crystal structure of SETDB1-TTD in complex with H3K9me2K14ac peptide (PDB: 6BHD.

We used a crystal soaking fragment screening approach to identify small-molecule ligands occupying the Kme and/or Kac pockets of SETDB1-TTD. Crystal structures revealed binding modes that could enable structure-guided hit optimization toward the development of SETDB1 antagonists.

## RESULTS AND DISCUSSIONS

### Hit Identification

In order to identify small molecule antagonists of SETDB1-TTD, we screened a fragment library of 252 compounds by soaking the apo crystals of SETDB1-TTD with pools of 3 compounds and identified a single fragment hit (**1**). Co-crystal structure of **1** with SETDB1-TTD revealed that it binds in the Kme pocket (Figure 2). However, fragment **1** did not show any HSQC shift at concentrations as high as 10 mM, indicating very weak binding.

**Figure 2.**
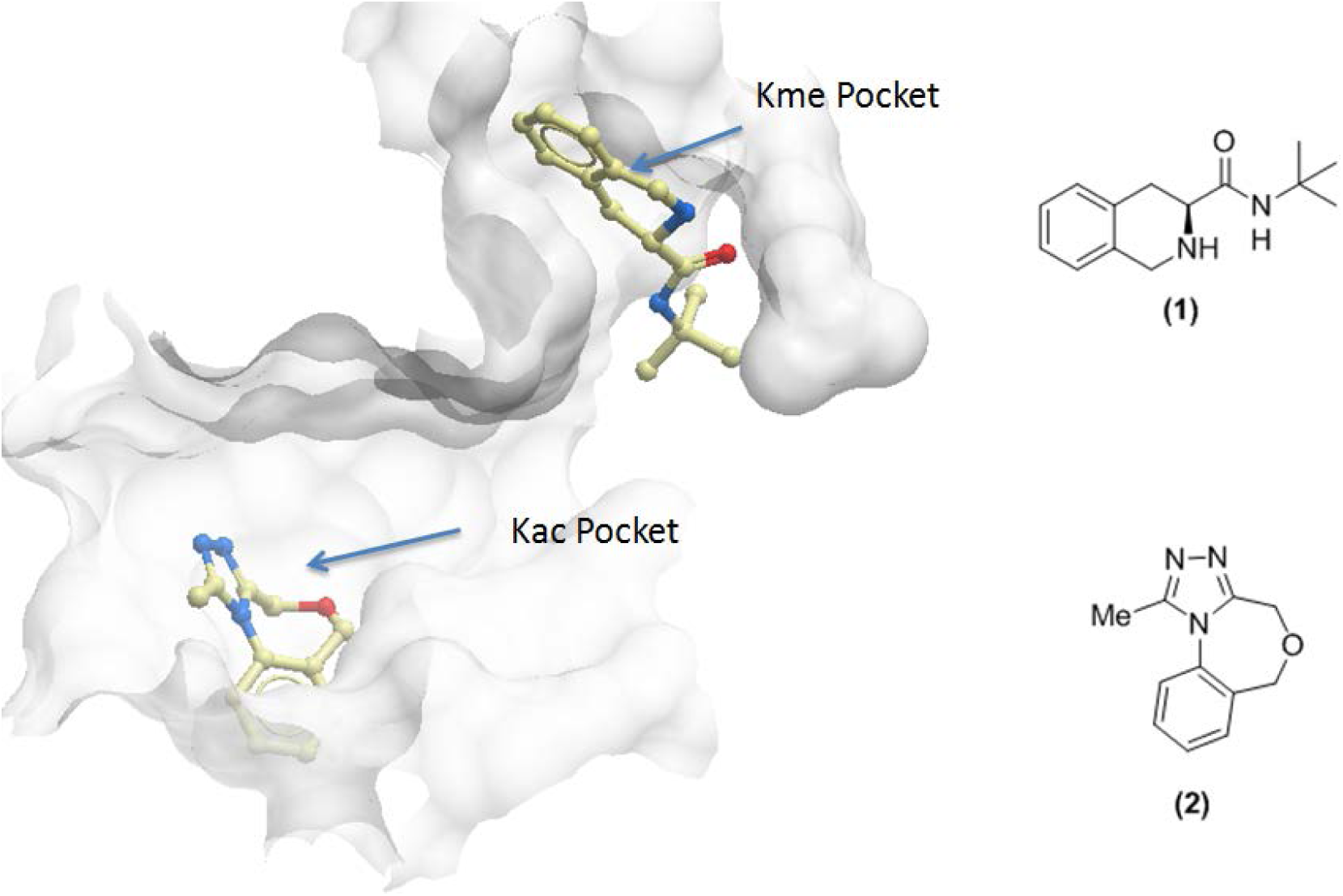
Superimposed crystal structures of the SETDB1 TTD in complex with **1** (PDB ID: 5KE2) and **2** (PDB ID: 6AU2) occupying the Kme and Kac pockets, respectively.

As a major part of the H3K9me2K14ac peptide occupies and forms hydrogen bonding interactions in and around the Kac pocket, fragments binding in the Kac pocket are likely to have higher ligand efficiency and can potentially be optimized to improve affinity. To test this hypothesis, we selected and soaked 5 constrained analogues of known bromodomain inhibitors^19^ synthesized in our laboratory (Supplementary Figure 1). Bromodomain inhibitors are known acetyl lysine mimics^20^ and could potentially bind to the Kac pocket of the SETDB1-TTD. Out of 5 fragments, one fragment (**2**) was found to bind to SETDB1-TTD occupying the TTD Kac pocket (Figure 2). Binding of **2** to SETDB1-TTD was also confirmed by HSQC NMR at 5mM (Supplementary Figure 2). In order to explore the SAR of these hits and assess the feasibility of improving their activity, a small set of analogs was synthesized and tested by HSQC NMR and/or by SPR.

### Chemistry

The tricyclic triazole **2** was prepared from commercially available 2-aminobenzyl alcohol (**11**), which was reacted with ethylchloroacetate **(12)** in the presence of sodium hydride to give the lactam. Treatment of the lactam with Lawesson’s reagent led to the corresponding thioamide **13**, which was subsequently condensed with acethydrazide **(14)** to give the desired tricycle **2** in 6% yield overall yield (Scheme 1).

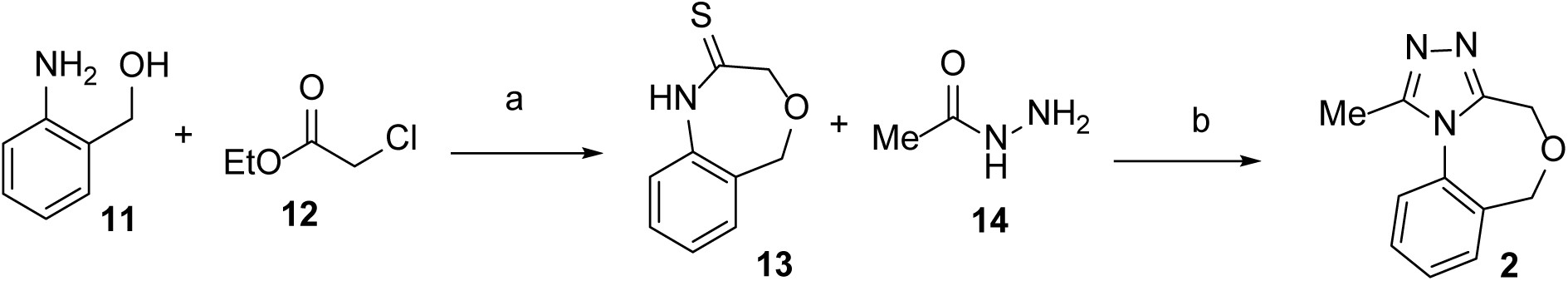
Scheme 1^a^. ^a^Reagents and conditions: (a) (1) NaH, Toluene, 80 °C; (2) Lawesson’s reagent, THF, 60 °C (b) ^n^BuOH, 100 °C, 6% overall yield.

*N*-Derivatized tricycle **4** was synthesized by the route outlined in Scheme 2. Diamine **15** was refluxed with ethylchloroacetate in toluene in the presence of triethylamine to obtain **16** as the major product. The crude product was selectively Boc-protected at the secondary amine to furnish **17**. The resulting material was carried directly through to intramolecular cyclization mediated by potassium *tert*-butoxide to give the lactam **18** in 35% isolated yield over three steps after recrystallization. Treatment of the cyclic amide with Lawesson’s reagent yielded thioamide **19**, which was then refluxed with acethydrazide for three days to furnish the triazole **20** in 80% yield after recrystallization. Finally, the Boc group was deprotected under acidic conditions and the resulting secondary amine was capped with acetyl chloride to give compound **4.**

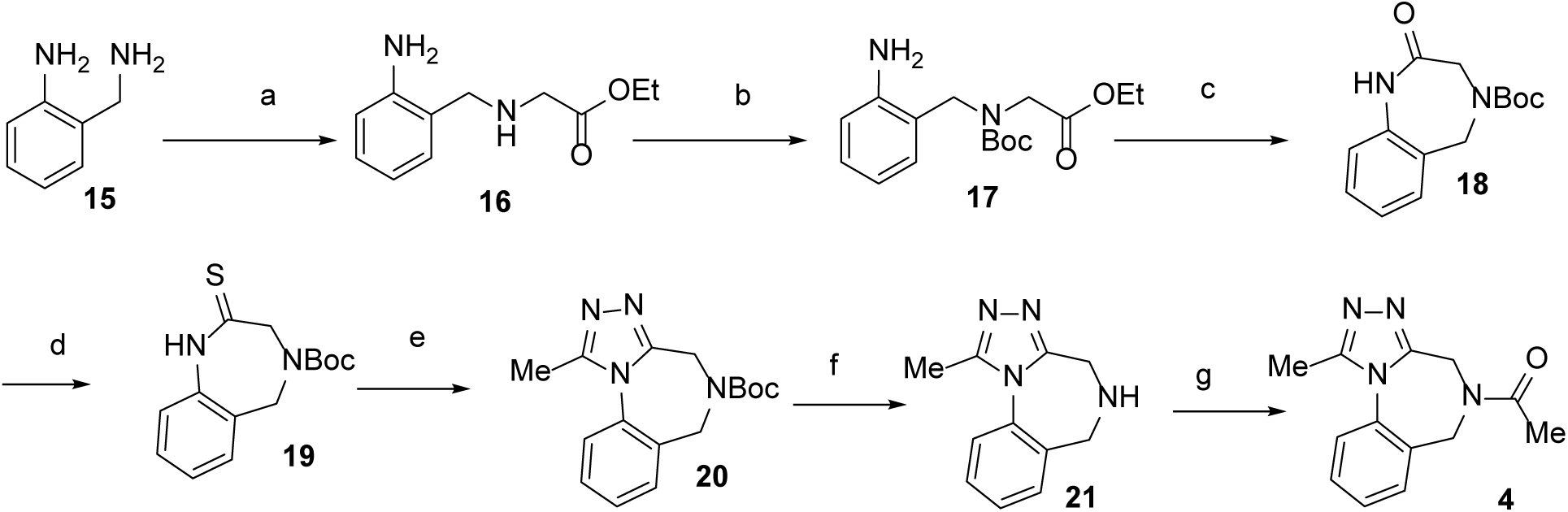
Scheme 2^a^. ^a^Reagents and conditions: (a) ethylchloroacetate, TEA, Toluene, 100 °C (b) Boc_2_O, cat.DMAP, THF, rt (c) KO^t^Bu, THF, 0 °C, 35% over 3 steps (d) Lawesson’s reagent, THF, 60 °C, 53% (e) acethydrazide, n-BuOH, 100 °C, 80% (f) 1.25 M HCl in MeOH, 50 °C, 97% (g) AcCl, TEA, DMF, rt, 54%

Triazoles **5-8** were prepared according to Scheme 3. Diamine **15** was treated with Boc anhydride followed by acetic anhydride to give the Boc-protected acetamide **25**, which was treated with Lawesson’s reagent to furnish thioamide **26**. Triazole **27** was obtained in moderate yield by treating **26** with acethydrazide under microwave irradiation. Deprotection of the Boc group was accomplished with hydrochloric acid in methanol and the free amine was treated with acetic anhydride or with methanesulfonyl chloride in the presence of base to yield the corresponding acetamide **5** and sulfonamide **7**, respectively. After the deprotection of triazole **27** under the same conditions, the free amine of the crude product was treated with triphosgene followed by methylamine to give **6**. The corresponding triazole carboxylic acid **8** was prepared by reacting **28** with acethydrazide.

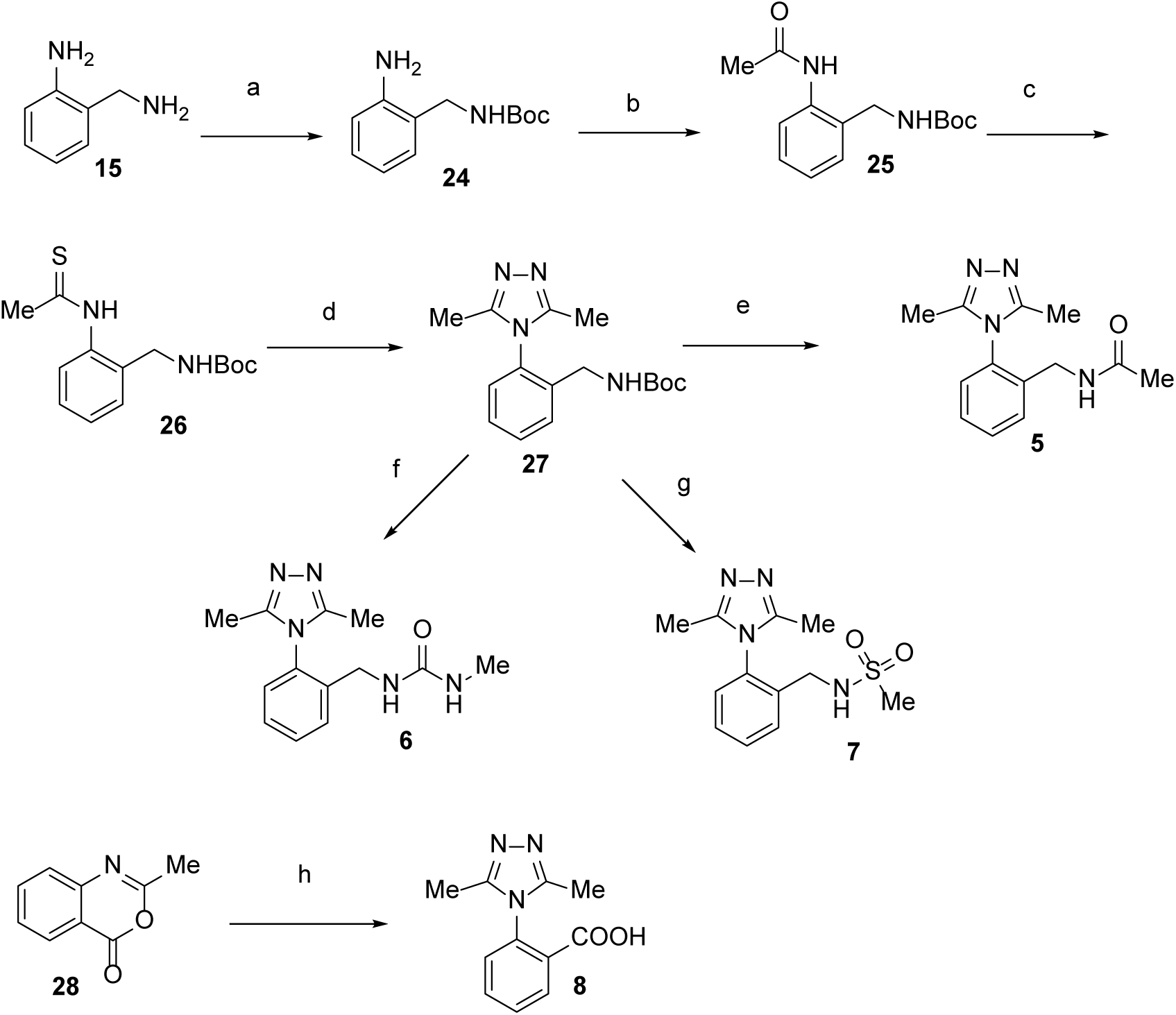
Scheme 3^a^. ^a^Reagents and conditions: (a) Boc_2_O, TEA, THF, rt, 99% (b) Ac_2_O, DCM, rt, 89% (c) Lawesson’s reagent, THF, 60 °C, 64% (d) acethydrazide, n-BuOH, 100 °C, 47% (e) (1) 1.25 M HCl in MeOH, 50 °C, 65%; (2) Ac_2_O, TEA, MeCN, rt, 65% (f) (1) 1.25 M HCl in MeOH, 50 °C; (2) triphosgene, TEA, DCM, rt; (3) MeNH_2_, MeOH, rt (g) (1) 1.25 M HCl in MeOH, 50 °C; (2) Methane sulfonyl chloride, TEA, MeCN, rt (h) acethydrazide, EtOH, reflux, 35%

*para*-Substituted triazole analogues **9a-e** were synthesized from intermediate **25** (Scheme 4). Halogenation of **25** with NBS and NCS gave **29a** and **29b**, respectively. Following the same protocol for converting **25** into **5** (Scheme 4), the corresponding halogenated intermediates **29a** and **29b** were converted to **9a** and **9b**. Palladium-catalyzed cyanation of **9a** gave **9c**, and Suzuki coupling of **9a** with the corresponding boronic acids gave **9d** and **9e**.

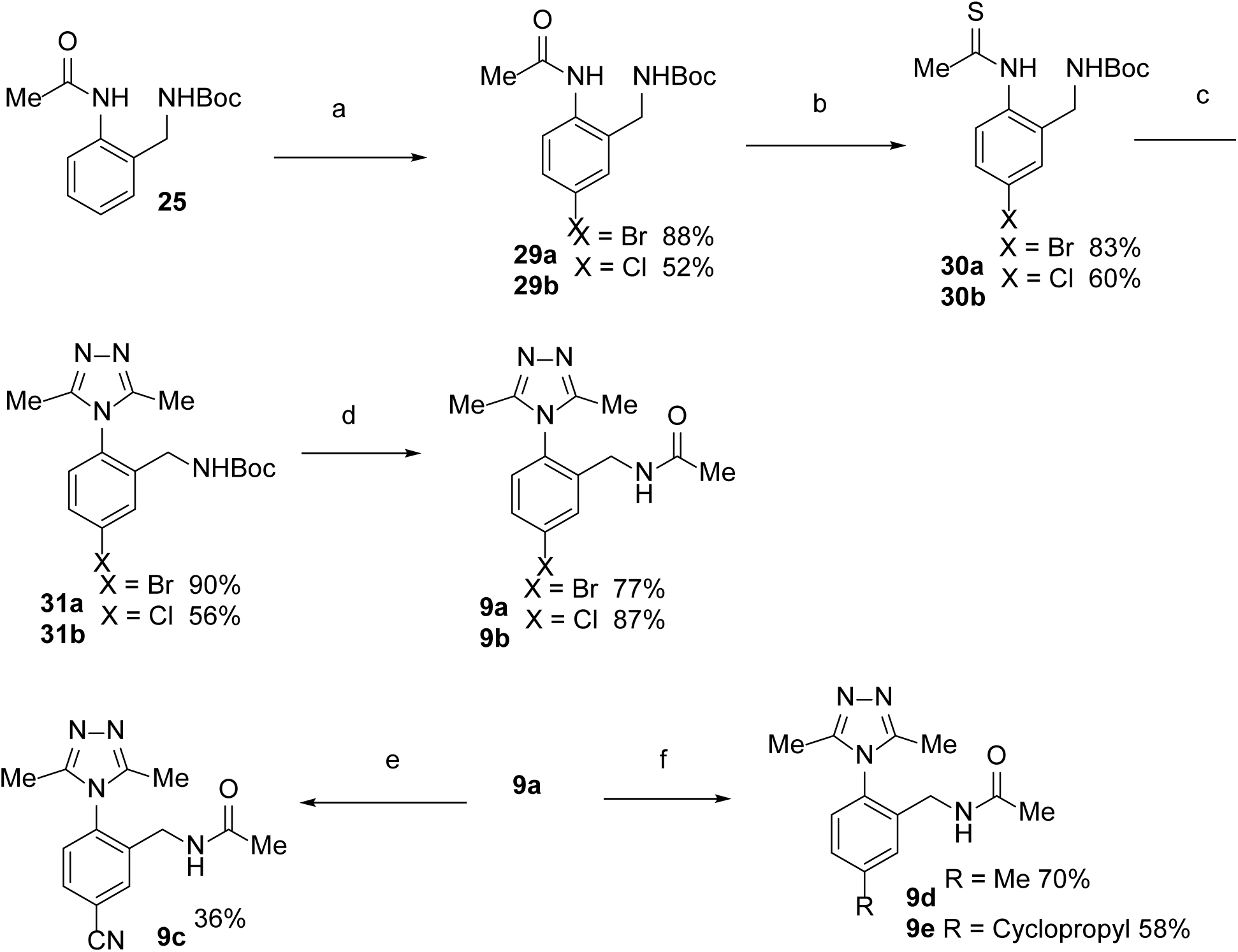
Scheme 4^a^. ^a^Reagents and conditions: (a) NBS or NCS, 10% triphenylphosphine sulfide, CHCl_3_, rt (b) Lawesson’s reagent, THF, 50 °C (c) acethydrazide, n-BuOH, 100 °C (d) (1) 1.25 M HCl in MeOH; (2) Ac_2_O, TEA, MeCN, rt (e) 10% Pd(OAc)_2_, K_4_Fe(CN)_6_, ^i^PrOH, DMF, 140 °C (f) 10% Pd(OAc)_2_, RB(OH)_2_, K_2_CO_3_, dioxane / water, 100 °C

Synthesis of the triazole fragment-linked molecule **10** began by condensing intermediate **28** with intermediate **32** to give the corresponding triazole carboxylic acid **33**. The alcohol **34,** which was obtained by the reduction of the carboxylic acid **33,** was converted into the corresponding phthalimide **35** under Mitsunobu conditions. Treatment of **35** with hydrazine to remove the phthalimide group followed by acetylation and deprotection of the Boc group gave intermediate **37**.

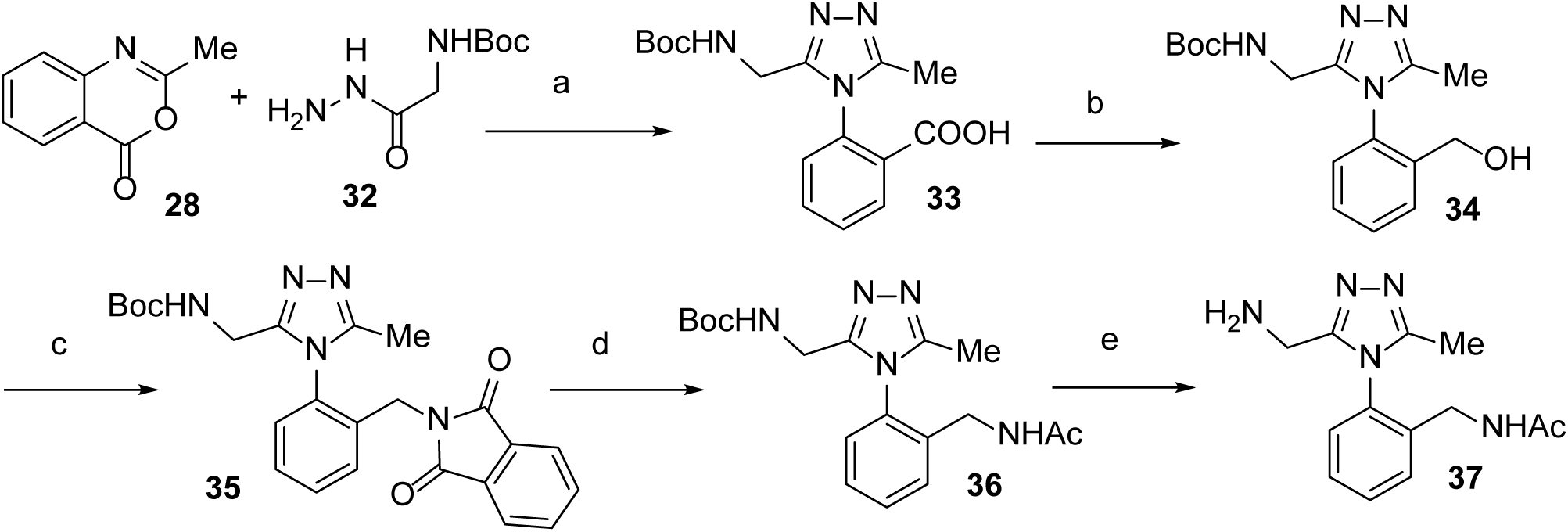
Scheme 5^a^. ^a^Reagents and conditions: (a) EtOH, reflux, 57% (b) LAH, THF, 54% (c) DIAD, PPh_3_, phthalimide, THF, rt, 96% (d) (1) hydrazine, MeOH, reflux; (2) N-acetyl-N-methoxyacetamide, MeOH, rt, 51% (e) TFA, 67%

The peptide intermediate **40** for the synthesis of **10** was synthesized from peptide **38**, which was in turn obtained using a standard 2-chlorotrityl resin protocol. Hydrogenolysis of the Cbz group of **38** followed by dimethylation *via* reductive amination of the lysine side chain with formaldehyde gave peptide **40** in 85% yield over two steps (Scheme 6). Coupling of the acid **40** with the amine **37** was followed by deprotection of Boc and ^t^Bu groups with TFA to give **10** (Scheme 7).

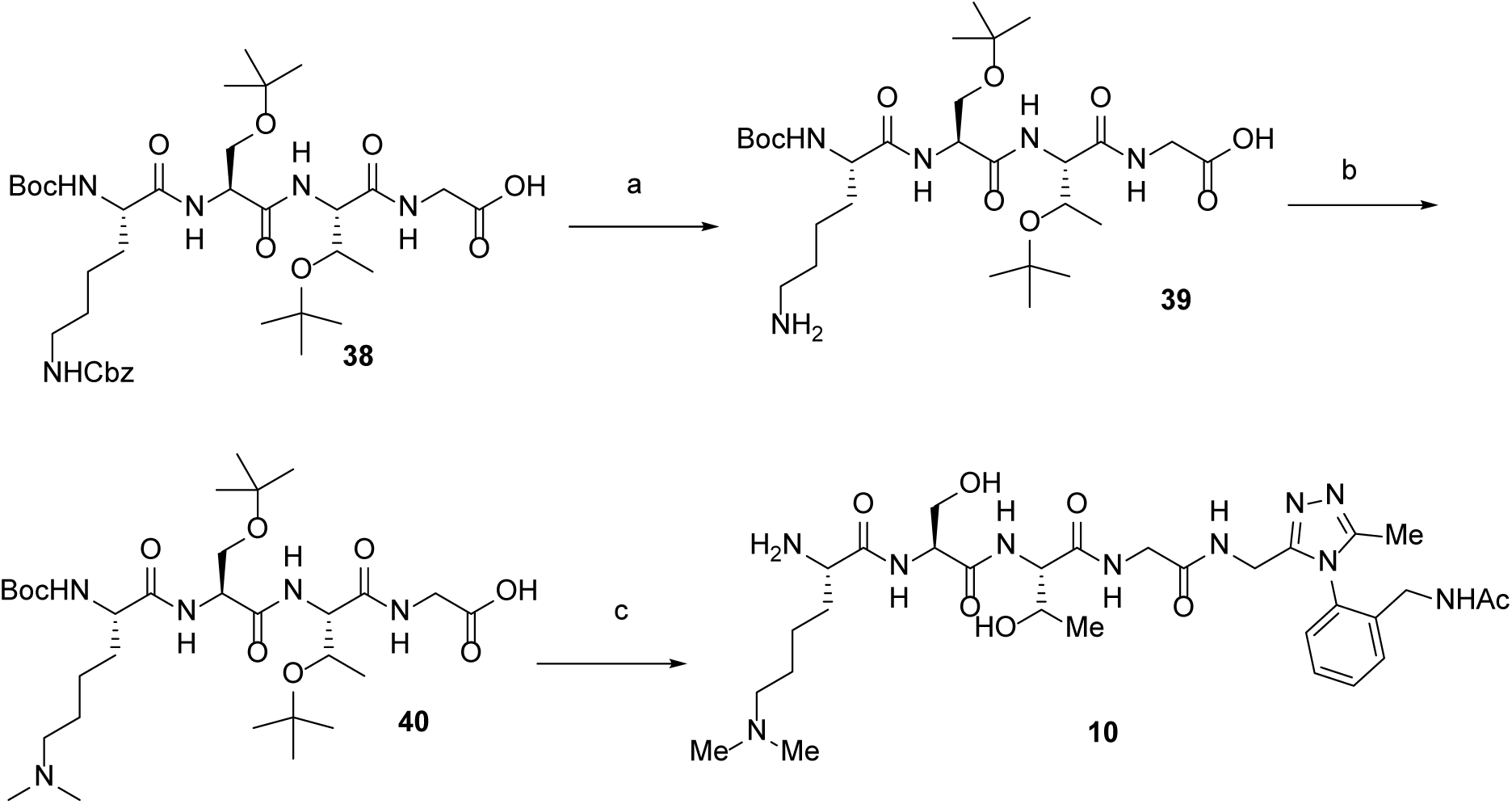
Scheme 6^a^. ^a^Reagents and conditions: (a) Pd/C, H_2_, AcOH, EtOH, quant. (b) HCHO, NaBH(OAc)_3_, MeOH, 85% (c) (1) EDC.HCl, DCM, rt; (2) TFA, DCM, rt, 30%

#### Biochemical evaluation and SAR of the fragments occupying Kme pocket

In the co-crystal structure of the fragment hit **1** with SETDB1-TTD, the phenyl ring of the tetrahydroisoquinoline core occupies the aromatic cage formed by W358, W363 and F379 and the carbonyl group makes hydrogen bonding with D382 (Figure 3a). However, **1** did not show any significant shift in the HSQC NMR of SETDB1-TTD at concentrations up to 10 mM, indicating that **1** interacts very weakly with the Kme pocket. Superimposed structures of SETDB1-TTD in complex with **1** and H3K9me2K14ac within the binding pocket (Figure 3b) indicate that the fragment could be grown towards the Kac pocket, which could potentially pick up additional interactions with the protein and thus improve affinity. We purchased a library of 19 amides (Supplementary figure 3) and synthesized 3 amides, extending fragment **1** in the desired direction. Ligand-protein interactions were assessed using HSQC NMR as a first pass screen. None of the 22 analogs tested showed HSQC NMR shift at 10 mM. However, we were able to obtain the co-crystal structure of **3** with SETDB1-TTD, wherein the tetrahydroisoquinoline core of **3** occupies the aromatic cage of the Kme pocket, and the amide extends towards the Kac pocket (Figure 3c). No electron density was observed for the furan group, indicating suboptimal interactions with the protein. As none of the analogs of **1** showed any significant activity, we turned our focus toward identifying fragments that occupy the Kac pocket.

**Figure 3.**
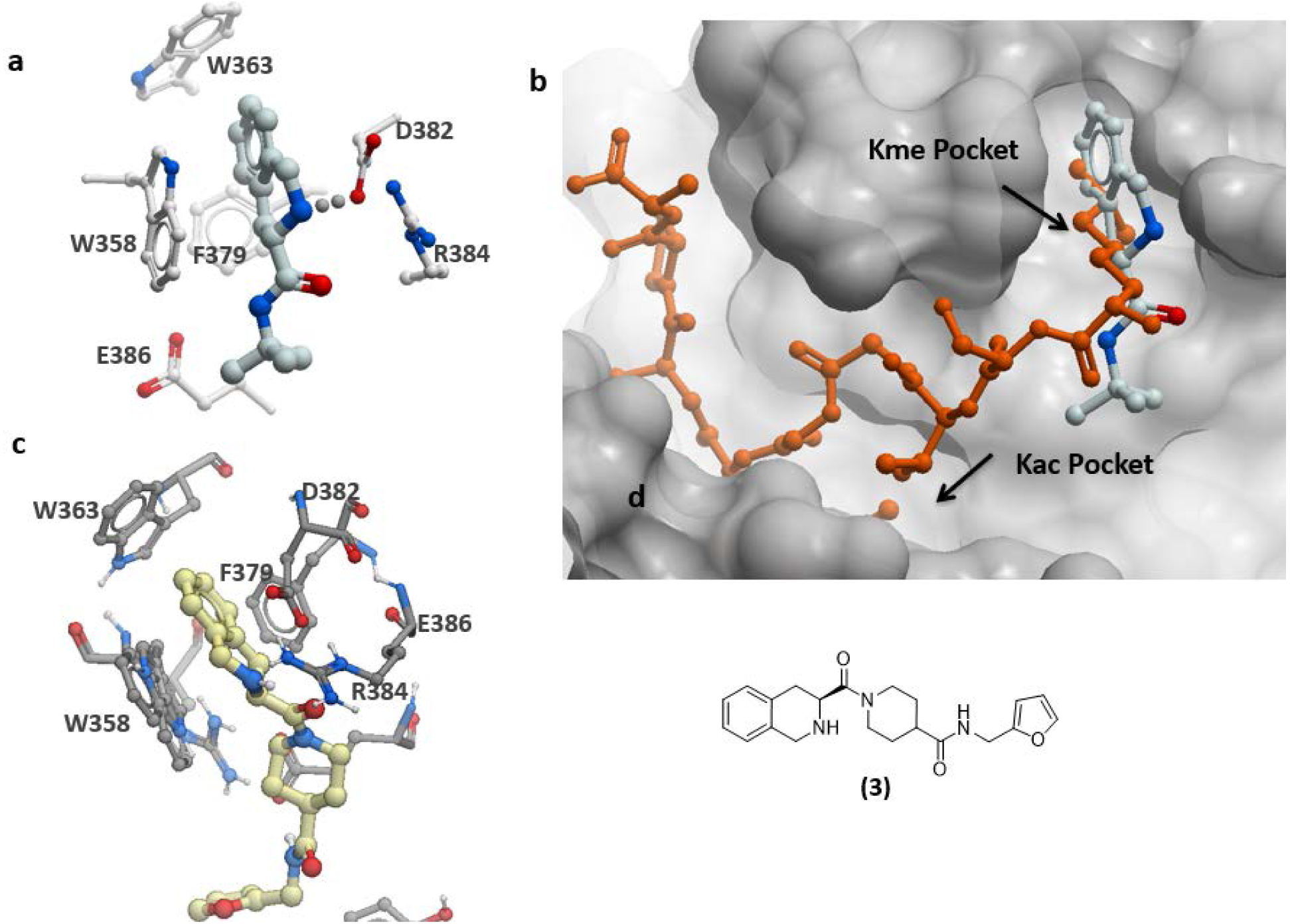
Crystal structure of SETDB1 TTD in complex with fragments. (a) Binding mode of **1** in the aromatic cage of Kme pocket. (b) Superimposed structures of **1** (grey) (PDB: 5KE2) with the co-crystal structure of H3K9me2K14ac (orange) (PDB: 6BHD). (c) Binding mode of **3** in the aromatic cage of the Kme pocket (PDB ID: 5KE3). (d) Chemical Structure of **3**.

#### Biochemical evaluation and SAR of fragments occupying the Kac pocket

In the co-crystal structure of SETDB1-TTD in complex with **2**, the triazole ring of **2** participates in π-stacking interactions with F332 and its phenyl ring is buried in the narrow pocket, forming hydrophobic interactions with I333, G300, F296, and C329. Both the nitrogen atom in the triazole ring and the oxygen atom in the seven membered ring form hydrogen bonds with two water molecules (Figure 4a). At 10 mM, **2** showed a significant HSQC NMR shift. Similarly, the simple acetamide derivative **4** showed a significant HSQC NMR shift (Supplementary Figure 2), but neither compound showed significant binding by ITC (not shown). In the H3K9me2K14ac co-crystal structure, the acetyl group of K14ac forms hydrogen bonding interactions with G300 and Y389, both of which are found in the vicinity of **2**. We therefore synthesized the corresponding open chain analogue **5** and obtained a co-crystal structure. As expected, the *N*-acetyl group of **5** formed hydrogen bonds with G300 and Y389 (Figure 4b). A slight shift in the scaffold binding pose compared with **2** produced an additional hydrogen bond between the trizole ring of **5** and the side-chain of R394, but this also leads to reduced interactions at the I333 hydrophobic cavity and loss of π-stacking with F332 (Figure 4c). Compound **5** showed a strong HSQC NMR shift at 5 mM (Supplementary Figure 2) and displayed K_D_ values in low millimolar ranges by ITC (K_d_=3.5 mM) (Figure 4d) and SPR (K_d_=3.5 mM) (Supplementary Figure 4).

**Figure 4.**
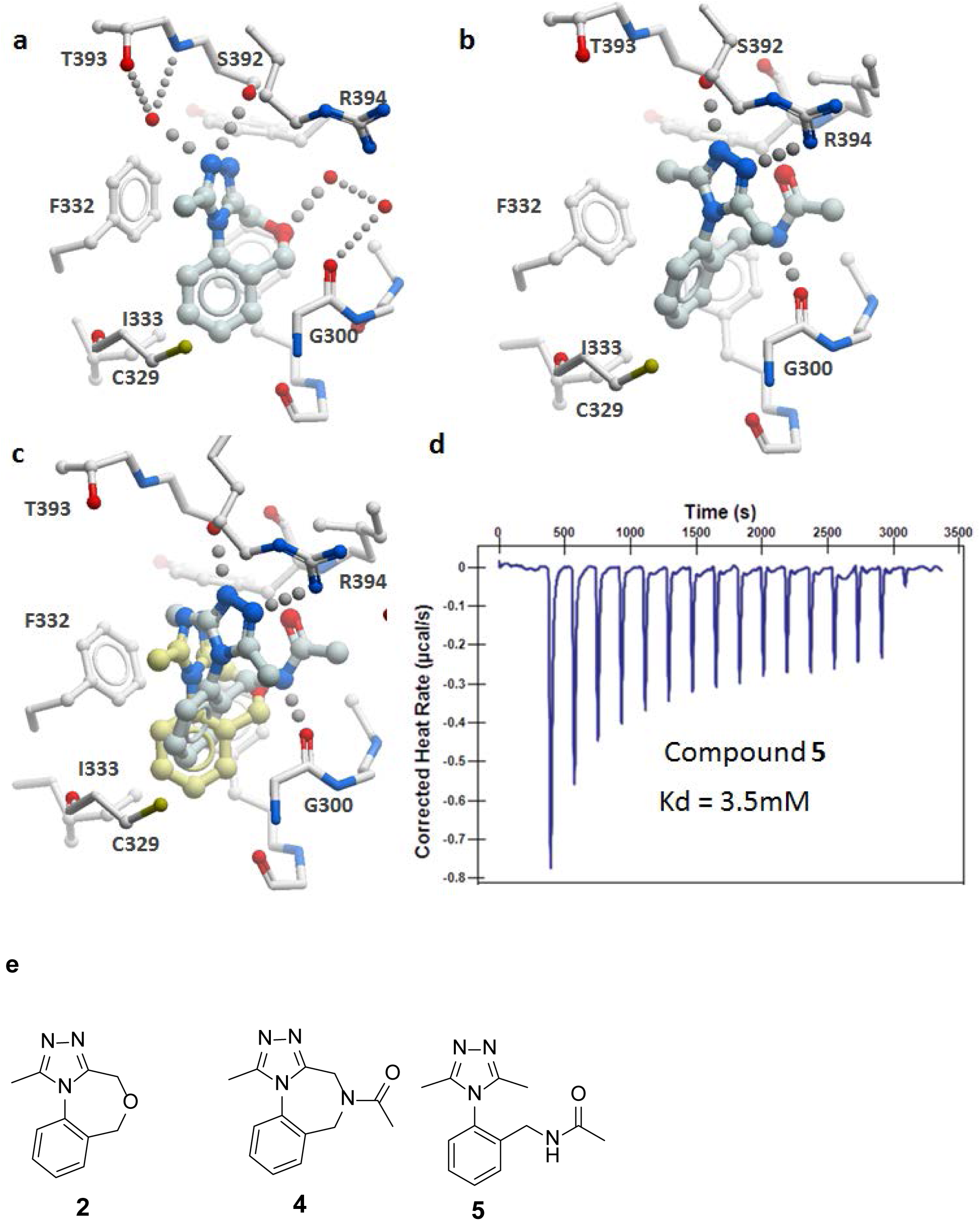
(a) Binding mode of **2** in the Kac pocket. (b) Binding mode of **5** in the Kac pocket (PDB ID: 6AU3). (c) Superimposed structures of **2** (yellow) with **5** (grey) (d) ITC data of **5**. (e) Chemical Structures of **2, 4** and **5**.

We also synthesized the corresponding urea **6**, sulfonamide **7**, and carboxylic acid **8** (Figure 5). All three compounds were predicted to form additional or alternate hydrogen bonds with the protein, however, only the sulfonamide analog **7** showed a strong HSQC NMR shift at 5 mM (Supplementary Figure 2) and affinity comparable to **5** by ITC (Figure 5) and SPR. (Supplementary Figure 4) (Kd = 4mM – 17mM in both assays).

**Figure 5.**
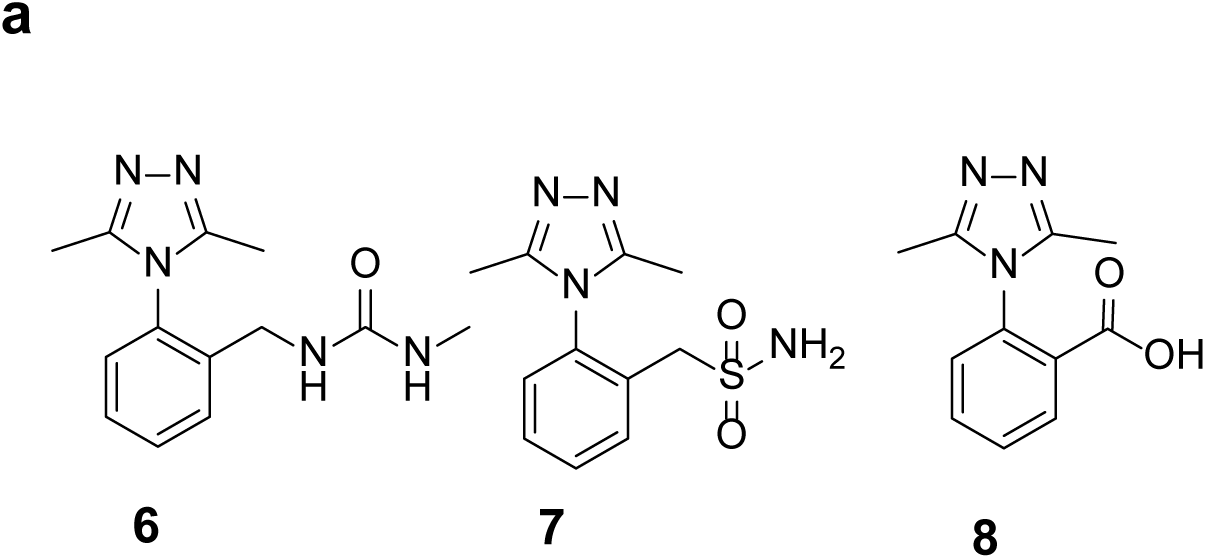
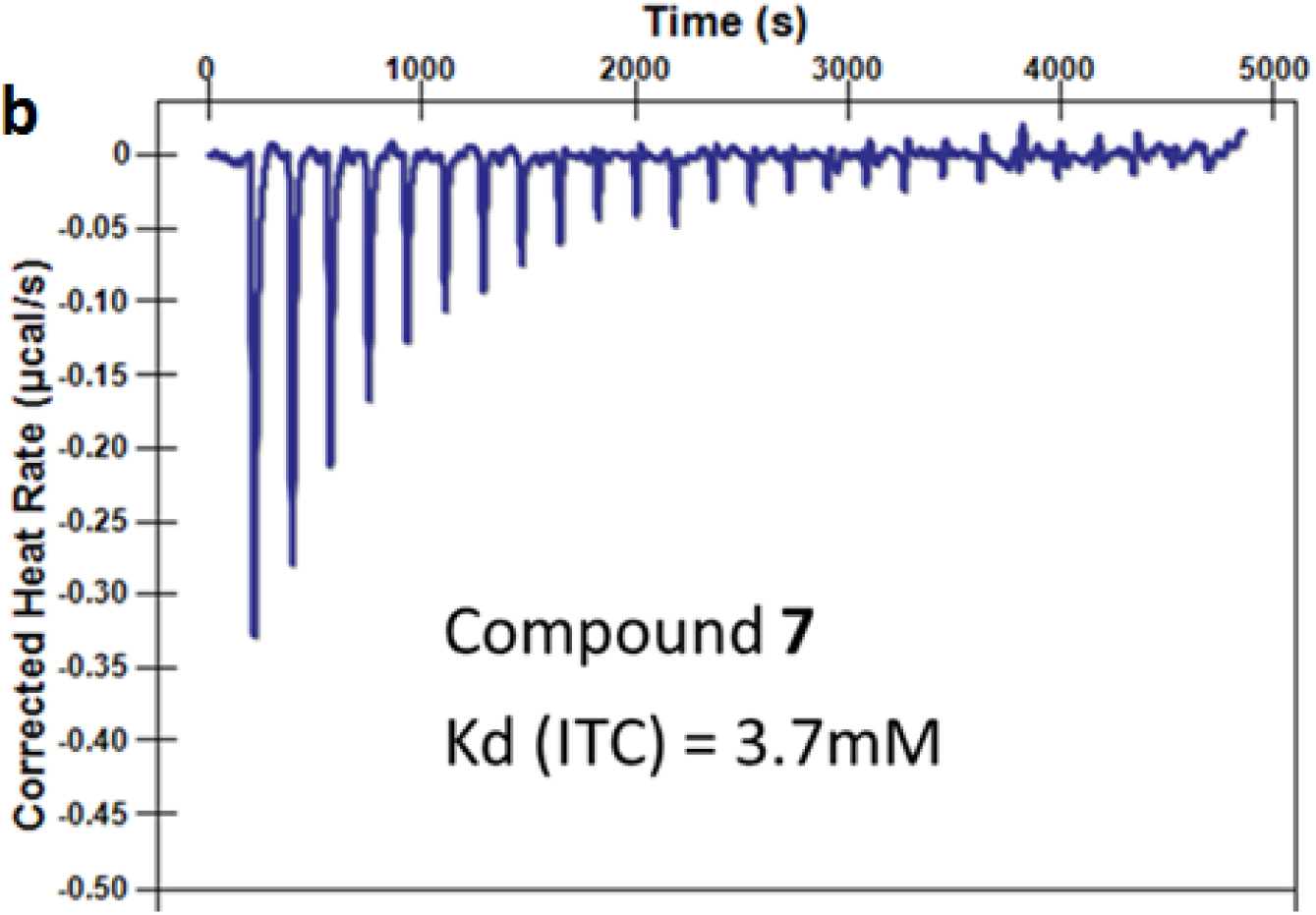
(a) Chemical Structures of **6-8**. (b) ITC data of **7**.

Despite forming a number of hydrogen bonding interactions with the protein, **5** showed only weak binding by ITC and SPR, which is likely due to incomplete occupation of the large pocket that is occupied by the native peptide H3K9me2K14ac. We therefore decided to grow the fragment **5** into the pocket occupied by the peptide. We initially focused on simple substitutions at the *para* position of the phenyl ring of **5** to rescue hydrophobic interactions observed between **2** and I333. However, none of the para-substituted analogues showed significant improvement of binding compared to **5** as measured by SPR (Table 1 and Supplementary Figure 4).

**Table 1.** SPR data of substituted analogues of **5.** Binding of compounds **5**, **9a**, **9b**, **9c** and **9d** to SETDB1 TTD was confirmed by SPR with K_d_ values mostly in single digit milimolar.

| Compound | R | K<sub>d</sub> (mM) |
| --- | --- | --- |
| **5** | H | 3.5 |
| **9a** | Br | 2.7 |
| **9b** | Cl | 2.5 |
| **9c** | CN | 10 |
| **9d** | Me | 7.7 |
| **9e** | Cyclopropyl | N/D |

N/D=Not determined (linear response)

Next, we linked fragment **5** with a portion of the H3K9me2K14ac peptide that occupies the desired Kme pocket and linker region to obtain compound **10.** The co-crystal structure indicates that **10** recapitulated the binding pose of **5** in the Kac pocket and the binding pose of H3K9me2K14ac in the Kme pocket and linker (Figure 6). Compound **10** showed a significant HSQC NMR shift, however displayed weak binding by SPR (SPR K_d_ = 1.27 mM), despite occupying and making interactions in the major part of the pocket occupied by the native peptide. This may be because the interactions in the cavities occupied by the native peptide, but not occupied by **10,** may be important for potency. This weak activity may reflect conformational strain associated with binding of the chemical series explored here. Structural dynamics of the TTD may also impact accessibility of the binding pocket.

**Figure 6.**
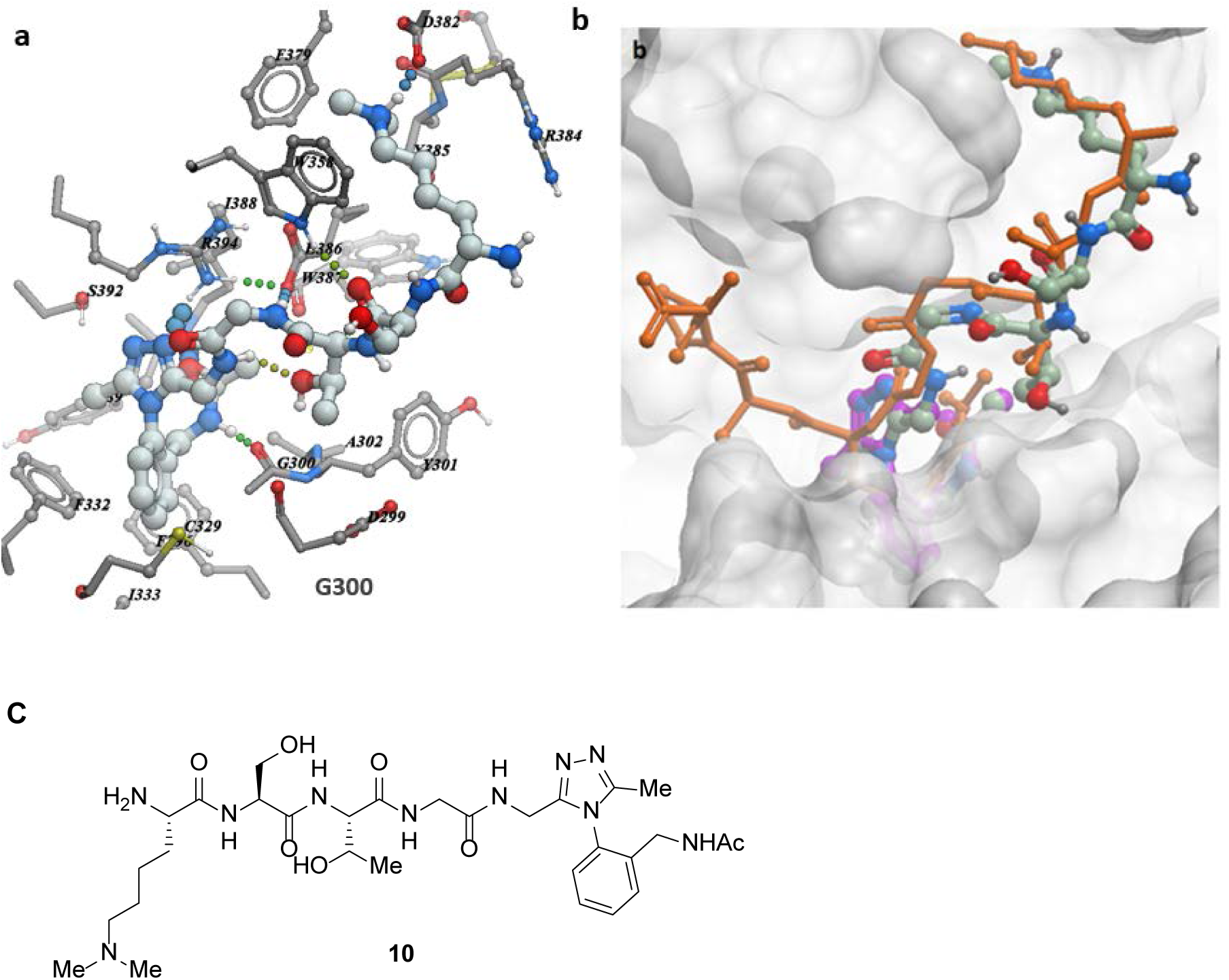
(a) Binding mode of **10** (PDB ID: 6BPI) (b) Overlay of the structures of **10** (green), **5** (purple) with the H3K9me2K14ac peptide (orange). (c) Chemical structure of **10.**

### Conclusion

We have identified the first fragment hits occupying the peptide binding pocket of SETDB1 TTD, which demonstrates its ligandability. Fragments occupying the Kac pocket form interactions with the protein. The most promising binding interactions were observed in compound 5, which included four hydrogen bonds and multiple hydrophobic interactions, however showed only weak binding by ITC and SPR. Compound **10**, a hybrid of fragment **5** and a histone peptide capable of spanning both Kme and Kac pockets, did not show significant binding, suggesting that cavities occupied by A15-R17 of the native peptide may be important for potency. Future compounds exploiting this portion of the binding site will help further define the structural chemistry of the SETDB1 TTD and map the road towards potent and selective SETDB1 chemical probes.

## EXPERMENTAL SECTION

### Protein expression and purification

The expression construct for N-terminal His6-tagged SETDB1 Tudor domain (residues 197 – 403) subcloned into pET28a-MHL vector was transformed into BL21(DE3)-pRARE2 cells. The cultures were grown in Terrific Broth medium (TB) in the presence of 50 µg/mL kanamycin and 34 µg/mL chloramphenicol at 37 °C. For NMR analyses, the cultures were grown in auto-induction M9 minimal medium containing 1.2 g/l 15N-ammonium chloride (Sigma 299251) as the sole nitrogen source using an auto-induction method. For SPR experiments, the expression construct was N-terminally biotinylated SETDB1 Tudor domain (residues 197 – 403) subcloned into pET28BIOH_LIC vector and growth was done as usual but supplemented with 1 mg/ml of biotin. The overexpression of SETDB1 Tudor was induced by addition of isopropyl-1-thio-D-galactopyranoside (IPTG), final concentration 0.5 mM, and incubated overnight at 16 °C. Next day, the cells were harvested by centrifugation at 12,227 × g (10 min, 4 °C). Details of the purification peocedures are provided in the supporting information.

### Protein crystallization

Purified SETDB1 was crystallized using hanging drop vapor diffusion method at 20 °C by mixing 1.5 µl of the protein solution at 5 mg/ml with 1.5 µl of the reservoir solution containing 25% PEG3350, 0.2 M LiSO4, 0.1 M Bis-Tris, pH 6.5. Matured SETDB1 crystals were transferred to the soaking solution containing 27% PEG3350, 0.2 M LiSO4, 0.1 M Bis-Tris, pH 6.5, 50 mM of the ligand and 0.5% DMSO. Soaking time varied from 24 to 48 hours.

### Biophysical assays

Isothermal titration calorimetry (ITC) for **5** was performed at 25 °C using Nano ITC instrument (TA Instruments, USA). The cell was loaded with 80 µM SETDB1 (Tudor domain) in 20 mM HEPES pH 7.5 and 250 mM NaCl. The syringe was loaded with 100 mM compound **5** dissolved in the same buffer (pH was checked and no DMSO was used). 25 injections of 2 µL volumes were applied at 180 s intervals. “Nano Analyze” software package (supplied by the manufacturer) was used for data analysis. The data was fit to a one binding site model.

ITC for compound **7** was performed using the same instrument and using similar conditions as for compound **5**. The cell was loaded with 98 µM SETDB1 in 20 mM HEPES pH 7.5 and 250 mM NaCl. The syringe was loaded with 50 mM compound **7** (It was not possible to prepare 100 mM solution due to limited solubility) dissolved in the same buffer as protein (again pH was checked and no DMSO was used).

NMR spectroscopy. 1H-15N HSQC-TROSY spectra were collected at 298 K on 500 MHz Bruker Avance spectrometer equipped with 1.7 mm micro cryoprobe. Experiments were performed at 5 mM ligand concentrations with 190 µM 15N labeled SETDB1 in 20 mM HEPES pH 7.5 and 250 mM NaCl, 2.5% DMSO, and 10% D2O. Negative controls contained 2.5% DMSO and no compound.

### General Chemistry

All solvents acetonitrile, dimethylformamide (DMF), tetrahydrofuran (THF), dichloromethane (DCM), and trifluoroacetic acid (TFA) were of reagent grade quality, purchased from commercial sources and used as received. Flash column chromatography was carried out using Silicycle 230- 400 mesh silica gel. Thin-layer chromatography (TLC) was performed on Macherey Nagel precoated glass-backed TLC plates (SIL G/UV254, 0.25 mm) and visualized using a UV lamp (254 nm) followed by KMnO_4_, ninhydrin or iodine stain. Reported R_f_ values are dependent on ambient humidity. NMR spectra were recorded at 25 °C on Bruker Advance III 400 MHz spectrometer, Varian Mercury 400, Agilent DD2 500, or Agilent DD2 600 instrument. Recorded shifts for protons are reported in parts per million (δ scale) downfield from tetramethylsilane and are referenced to residual protons in the NMR solvents (CDCl_3_: 7.26 or CD_3_CN: δ 1.94, centre line). Chemical shifts for carbon resonances are reported in parts per million (δ scale) downfield from tetramethylsilane and are referenced to the carbon resonances of the solvent (CDCl_3_: δ 77.16 or CD_3_CN: δ 1.32, centre line). Data are represented as follows: chemical shift δ in ppm, multiplicity (s singlet, d doublet, t triplet, q quartet, m multiplet, br broad), coupling constant *J* in Hz and integration. High resolution mass spectra were obtained on a VG 70-250S (double focusing) mass spectrometer at 70 eV or on an ABI/Sciex Qstar mass spectrometer with ESI source, MS/MS and accurate mass capabilities or on JEOL AccuTOF-DART instrument.

All the test compounds synthesized showed ≥95% pure as determined by HPLC/MS. Low-resolution mass spectra (ESI) were collected on an Agilent Technologies 1200 series HPLC paired to a 6130 Mass Spectrometer. Compounds were resolved on Phenomenex’s Kinetex 2.6u C18 50×4.6mm column at room temperature with a flow of 1 mL/min. The gradient consisted of eluents A (0.1% formic acid in double distilled water) and B (0.1% formic acid in HPLC-grade acetonitrile). Method A: A linear gradient starting from 5% of B to 95% over 15 min at a flow rate of 1.0 mL/min. Method B: A linear gradient starting from 5% of B to 95% over 4.00 min at a flow rate of 1.0 mL/min. Stays constant at 95% for 1 min and then returns to 5% over 0.5 min. Method C: Stays constant at 5% of B for 1.00 min at a flow rate of 1.0 mL/min, followed by a linear gradient to 95% over 8.0 min. Stays constant at 95% of B for 1.00 min and then returns to 5% B over 1.00 min. Compound 2 (Method A); Compounds 4, 9c, 9d (Method B); Compounds 6-8, 9a, 9b, 9e (Method C).

### Synthetic procedures. General procedure A: synthesis of thioamides

Amide (1 equiv) was dissolved in THF (0.1 M) and Lawesson’s reagent (0.6 – 0.75 equiv) was added. The reaction was heated to 60 °C for 2 – 4 hours until complete by TLC analysis. Excess silica gel was added to the stirred solution and was allowed to cool to room temperature while stirring. The solvent was evaporated *in vacuo* and the product dry loaded onto silica gel. The silica-loaded crude product was loaded onto a silica column and the product carefully eluted (0% to 30% EtOAc in hexanes) as a yellow band following the initial dark byproduct band.

### General procedure B: Boc deprotection and acetylation of amines

The Boc-protected amine (1 equiv) was dissolved in 1.25 M HCl in methanol (15 equiv) and the solution was heated to 50 °C overnight. The reaction achieved full conversion as gauged by LC/MS analysis and the solvent was evaporated under reduced pressure. The residue was dissolved in acetonitrile (0.2 M) and Amberlite IRA-67 was added to free-base the amine followed by addition of acetic anhydride (1.5 equiv). The reaction was allowed to stir for 6 – 12 hours and after removal of the solvent, crude products were purified by reverse phase or normal phase column chromatography.

### General procedure C: Synthesis of triazoles from thioamides

Thioamide (1 equiv) was added to a microwave vial followed by acethydrazide or formic hydrazide (4 – 5 equiv). n-Butanol (0.5 M) was added and the vial capped and heated at 100 °C under normal microwave irradiation for 24 – 36 hours until the starting material was consumed as gauged by LC/MS analysis. The solvent was evaporated *in vacuo* and the hydrazide sublimed from the reaction mixture under high vacuum at 100 °C. The crude product was subsequently purified by reverse phase column chromatography or by recrystallization.

General procedure D: Synthesis of para-halogenated N-acylanilines. N-acylaniline (1 equiv), NBS/NCS (2 equiv) and triphenylphosphine sulfide (0.1 equiv) were dissolved in chloroform (0.05 M) and stirred for 1 (NBS) to 2 (NCS) days until the starting material was fully consumed. The solvent was evaporated in vacuo and the crude product was redissolved in EtOAc, washed subsequently with water, saturated sodium thiosulfate, water and brine and dried over sodium sulfate. The crude product was then purified by column chromatography.

### 1-methyl-4*H*,6*H*-benzo[*e*][1,2,4]triazolo[3,4-*c*][1,4]oxazepine (2)

2-aminobenzylalcohol **(11)** (500 mg, 4.06 mmol) was combined with sodium hydride (195 mg, 4.87 mmol) in toluene (8.1 mL) and added with ethylchloroacetate **(12)** (0.435 mL, 4.06 mmol). The solution was heated to 80 °C for 1 hour, cooled to room temperature, diluted with EtOAc and worked up with 10% aqueous HCl. The organic layer was washed with water, brine, and dried over sodium sulfate. After passing through silica column the crude lactam (74.2 mg, ~11%, pale yellow solid (*R_f_* (1:1 EtOAc:hexanes): 0.33) was carried through to the next step. Crude lactam (73.7 mg, 0.45 mmol) was dissolved in toluene (4.52 mL) and Lawesson’s reagent (183.1 mg, 0.45 mmol) was added. The solution was heated to 60 °C for 5 hours, concentrated and passed through a silica gel plug (*R_f_* (2:5 EtOAc:hexanes): 0.64). The crude thioamide product **13** (0.092 g, 0.52 mmol) was mixed in *n*-butanol (2.2 mL) and acethydrazide **(14)** (76.3 mg, 1.03 mmol) was added. The solution was heated to 120 °C for four days, the solvent removed *in vacuo* and the crude product was purified by column chromatography. Yield: 48.0 mg, 53%, pale yellow crystals. *R_f_* (1:19 MeOH:EA): 0.1.^1^H NMR (500 MHz, CDCl_3_) *δ* 7.60 (d, *J* = 7.9, 7.4, 1.7 Hz, 1H), 7.55 (ddd, *J* = 7.6, 1.8, 0.5 Hz, 1H), 7.49 (dt, *J* = 7.5, 1.2 Hz, 1H), 7.39 (dd, *J* = 8.0, 0.9 Hz, 1H), 4.66 (s, 2H), 4.46 (s, 2H), 2.62 (s, 3H). ^13^C NMR (126 MHz, CDCl_3_) δ 151.8, 150.9, 134.4, 131.9, 131.1, 130.4, 129.3, 122.6, 67.1, 59.2, 11.9. IR (neat) *ṽ* (cm^-1^): 2928, 2864, 1526, 1495, 1462, 1423, 1380, 1087, 878, 770. HRMS (DART) *m/z*: [M + H]^+^ calcd for C_11_H_12_N_3_O 202.0980, found 202.0980.

### *tert*-butyl 2-oxo-1,2,3,5-tetrahydro-4*H*-benzo[*e*][1,4]diazepine-4-carboxylate (18)

In a flame dried round-bottom flask, 2-aminobenzylamine (10 g, 82 mmol) was dissolved in toluene (410 mL, 0.2 M). Triethylamine (12.6 mL, 90 mmol) and ethylchloroacetate (9.2 mL, 86 mmol) were added and the solution was stirred under reflux at 110 °C for 22 hours, until complete as indicated by TLC and LC/MS. The reaction mixture was cooled to room temperature and water (410 mL) was added. The organic layer was separated and washed with water, followed by brine, dried over sodium sulfate and concentrated to yield ethyl (2-aminobenzyl)glycinate **(16)** which was used in the subsequent step without further purification. Yield: 15.8 g, 93%, brown/orange oil. *R_f_* (EtOAc): 0.51. ^1^H NMR (300 MHz, CDCl_3_) δ 7.27 – 7.15 (m, 1H), 7.13 – 7.07 (m, 1H), 7.06 – 7.00 (m, 1H), 6.69 – 6.67 (m, 1H), 4.24 – 4.17 (m, 2H), 3.79 (s, 2H), 3.40 (s, 2H), 1.31 – 1.27 (m, 3H). ^13^C NMR (100 MHz, CDCl_3_) δ 172.7, 146.9, 130.3, 128.7, 123.2, 117.8, 115.8, 60.9, 52.1, 49.8, 14.3. IR (neat) *ṽ* (cm^-1^): 3436, 3342, 2976, 1730, 1617, 1495, 1459, 1192, 1131, 749. The crude product **16** (15.8 g, 75.9 mmol) was dissolved in THF (760 mL) followed by the addition of Boc_2_O (21.8 mL, 94.8 mmol) and a catalytic amount of DMAP. The solution was allowed to stir for 20 hours until reaction was complete by TLC analysis. The THF was removed under reduced pressure, and the residue redissolved in EtOAc, washed with saturated sodium bicarbonate, brine, and dried over sodium sulfate. The volatiles were removed *in vacuo* and the crude product **17** (*R_f_* (1:3 EtOAc:hexanes): 0.7) was dissolved in THF (760 mL) and cooled to 0 °C. Potassium *tert*-butoxide (8.94 g, 79.7 mmol) was added portionwise until the reaction was complete by TLC. The solvent was evaporated, the crude redissolved in EtOAc, washed with saturated sodium bicarbonate, then brine, then dried over sodium sulfate. Compound **18** was purified by recrystallization from hot EtOAc. Yield: 7.6 g, 35% (3 steps), white solid. *R_f_* (1:3 EtOAc:hexanes): 0.2. 1:1 Mixture of rotamers. ^1^H NMR integrations adjusted to reflect whole number protons of each rotamer in solution. ^1^H NMR (500 MHz, CDCl_3_) δ 8.10 (br. s, 1H), 7.92 (br. s, 1H), 7.37 – 7.25 (m, 3H), 7.22 – 7.16 (m, 1H), 7.16 – 7.03 (m, 3H), 6.92 (d, *J* = 7.9 Hz, 2H), 4.55 (br. s, 2H), 4.48 (br. s, 2H), 4.42 (br. s, 2H), 4.29 (br. s, 2H), 1.44 (br. s, 9H), 1.37 (br. s, 9H). ^13^C NMR (126 MHz, CDCl_3_) *δ* 171.1, 170.3, 154.4, 153.9, 136.3, 130.2, 129.6, 128.8, 128.6, 127.9, 125.0, 123.9, 120.3, 119.9, 81.1, 80.8, 51.6, 51.2, 49.9, 48.9, 28.2. IR (neat) *ṽ* (cm^-1^): 3215, 3061, 3003, 1681, 1661, 1590, 1385, 1366, 1166, 1132, 752. HRMS (DART) *m/z*: [M + H]^+^ calcd for C_14_H_22_N_3_O_3_ 263.1396, found 263.1399.

### *tert*-butyl 2-thioxo-1,2,3,5-tetrahydro-4*H*-benzo[*e*][1,4]diazepine-4-carboxylate (19)

Synthesized according to general procedure A. Scale: 7.62 mmol. Yield: 1.13 g, 53%, pale yellow solid. *R_f_* (EtOAc): 0.93. ^1^H NMR (300 MHz, CDCl_3_) δ 9.50 (s, 1H), 7.34 (dt, *J* = 7.6, 1.6 Hz, 2H), 7.24 (d, *J* = 7.5 Hz, 1H), 7.04 – 6.99 (m, 1H), 4.53 (m, 4H), 1.46 (s, 9H). ^13^C NMR (100 MHz, CDCl_3_) *δ* 200.6, 154.3, 130.8, 129.3, 127.2, 121.1, 81.3, 56.2, 48.5, 28.5. IR (neat) *ṽ* (cm^-1^): 3239, 2982, 1659, 1488, 1417, 1362, 1153, 1119, 753. HRMS (ESI) *m/z*: [M + H]^+^ calcd for C_14_H_19_N_2_O_2_S 279.1167, found 279.1167.

### *tert*-butyl 1-methyl-4*H*-benzo[*f*][1,2,4]triazolo[4,3-*a*][1,4]diazepine-5(6*H*)-carboxylate (20)

Thioamide **19** (0.6 g, 2.16 mmol) and acethydrazide (0.24 g, 3.23 mmol) were dissolved in *n-*butanol, and stirred at 110 °C for 3 days. The reaction mixture was concentrated and acethydrazide was sublimed from the reaction mixture at 110 °C under high vacuum. The crude was recrystallized from hot EtOAc followed by recrystallization from methanol/diethylether. Yield: 516 mg, 80%, white solid. ^1^H NMR (500 MHz, CDCl_3_) δ 7.63 – 7.51 (m, 2H), 7.50 – 7.43 (m, 1H), 7.36 (d, *J* = 7.9 Hz, 1H), 4.58 (br. s, 2H), 4.37 (br. s, 2H), 2.58 (s, 3H), 1.50 (br. s, 9H). ^13^C NMR (126 MHz, CDCl_3_) δ 153.9, 151.2, 150.2, 133.5, 131.7, 130.9, 129.6, 129.4, 123.0, 81.1, 45.6, 40.8, 28.4, 11.8. IR (neat) *ṽ* (cm^-1^): 2979, 1693, 1497, 1384, 1259, 1156, 862, 762. HRMS (DART) *m/z*: [M + H]^+^ calcd for C_16_H_21_N_4_O_2_ 301.1665, found 301.1664.

### 1-(1-methyl-4*H*-benzo[*f*][1,2,4]triazolo[4,3-*a*][1,4]diazepin-5(6*H*)-yl)ethan-1-one (4)

Synthesized according to general procedure B. Scale: 0.33 mmol. Yield: 43.6 mg, 54%, white solid. *R_f_* (1:19 MeCN:DCM): 0.26. 1:2 mixture of rotamers. Methyl integral of 3 corresponds to major rotamer. ^1^H NMR (500 MHz, CDCl_3_) δ 7.66 – 7.34 (m, 6H), 4.74 –4.44 (m, 5H), 4.36 (s, 1H), 2.61 (s, 3H), 2.59 (s, 1.5H), 2.28 (s, 1.5H), 2.20 (s, 3H). ^13^C NMR (126 MHz, CDCl_3_) δ 169.4, 168.9, 151.5, 151.5, 150.4, 149.7, 133.7, 133.5, 132.1, 130.7, 130.6, 130.5, 130.3, 123.0, 129.7, 129.6, 123.7, 123.1, 48.9, 44.4, 42.5, 40.3, 22.2, 22.1, 12.0, 11.9. IR (neat) *ṽ* (cm^-1^): 3493, 3003, 1627, 1608, 1500, 1417, 1235, 1220, 776. HRMS (DART) *m/z*: [M + H]^+^ calcd for C_13_H_15_N_4_O 243.1246, found 243.1236.

### *tert*-butyl (2-acetamidobenzyl)carbamate (25)

2-Aminobenzylamine (3.0 g, 24.6 mmol) was dissolved in 50 mL of dry THF. Triethylamine (3.8 mL, 27.2 mmol) was added to the solution followed by Boc_2_O (5.4 g, 24.6 mmol). The mixture was stirred overnight at room temperature under a N_2_ atmosphere. The reaction mixture was poured into a 10% citric acid solution (25 mL) and extracted with EtOAc (2 x 50 mL). The combined organics were washed with brine (25 mL), dried over MgSO_4_, filtered and concentrated to give **24** as a white solid in 99% yield (5.41 g, 24.3 mmol). In a flame dried round-bottom flask, *tert*-butyl(2-aminobenzyl)carbamate **24** (1.5 g, 6.75 mmol) was dissolved in DCM (13.5 mL) and acetic anhydride (0.7 mL, 7.43 mmol) was added to the solution at 0 °C and allowed to warm to room temperature. After 2 hours, the reaction reached completion by TLC. The reaction mixture was diluted with dichloromethane and quenched with saturated sodium bicarbonate solution. The solution was washed with brine and dried over magnesium sulfate. The solvent was removed *in vacuo* and the product purified by column chromatography. Yield: 1.58 g, 89%, white solid. *R_f_* (1:1 EtOAc:Hexanes): 0.52. ^1^H NMR (400 MHz, CDCl_3_) δ 9.41 (s, 1H), 8.24 (d, *J* = 8.2 Hz, 1H), 7.31 (ddd, *J* = 8.5, 7.4, 1.7 Hz, 1H), 7.14 – 7.11 (m, 1H), 7.03 (dt, *J* = 7.4, 1.3 Hz, 1H), 5.07 (s, 1H), 4.27 (d, *J* = 6.8 Hz, 2H), 2.24 (s, 3H), 1.45 (s, 9H). ^13^C NMR (125 MHz, CDCl_3_) δ 169.5, 157.4, 137.2, 130.4, 129.2, 186.2, 124.0, 122.4, 80.9, 42.0, 28.5, 24.6. IR (neat) *ṽ* (cm^-1^): 3339, 3288, 2982, 1689, 1670, 1527, 1489, 1287, 1255, 1164, 748. HRMS (ESI) *m/z*: [M + Na]^+^ calcd for C_14_H_20_N_2_NaO_3_ 287.1366, found 287.1365.

### *tert*-butyl (2-ethanethioamidobenzyl)carbamate (26)

Synthesized using general procedure A. Scale: 2.74 mmol. Yield: 494 mg, 64%, pale yellow solid. *R_f_* (1:1 EtOAc:Hexanes): 0.8. ^1^H NMR (400 MHz, CDCl_3_) δ 8.17 (d, *J* = 8.1 Hz, 1H), 7.39 – 7.35 (m, 1H), 7.25 (m, 1H), 7.23 – 7.21 (m, 1H), 4.18 (s, 2H), 2.82 (s, 3H), 1.42 (s, 9H). ^13^C NMR (125 MHz, CDCl_3_) δ 210.5, 157.6, 137.7, 132.3, 130.5, 128.3, 127.2, 126.7, 81.1, 41.2, 35.5, 28.5. IR (neat) *ṽ* (cm^-1^): 3273, 2928, 1669, 1609, 1530, 1365, 1288, 1151, 1139, 851, 743, 701. HRMS (ESI) *m/z*: [M + H]^+^ calcd for C_14_H_21_N_2_O_2_S 281.1318, found 281.1330.

### *N*-(2-(3,5-dimethyl-4*H*-1,2,4-triazol-4-yl)benzyl)acetamide (5)

Synthesized according to general procedure B. Scale: 0.314 mmol. Yield: 50 mg, 65%, white solid. *R_f_* (1:9 MeOH:EtOAc): 0.2. ^1^H NMR (500 MHz, CDCl_3_) δ 7.70 – 7.64 (m, 1H), 7.53 (t, *J* = 7.6, 1H), 7.46 – 7.42 (m, 1H), 7.09 – 7.06 (m, 1H), 4.18 – 4.08 (m, 2H), 2.20 (s, 6H), 2.00 (s, 3H). ^13^C NMR (126 MHz, CDCl_3_) δ 170.4, 151.5, 136.3, 132.2, 130.9, 130.6, 129.1, 127.8, 39.7, 22.9, 11.0. IR (neat) *ṽ* (cm^-1^): 3362, 3253, 2992, 1687, 1666, 1542, 1512, 1279, 1270. HRMS (ESI) *m/z*: [M + H]^+^ calcd for C_13_H_17_N_4_O 245.1397, found 245.1397.

### 1-(2-(3,5-dimethyl-4*H*-1,2,4-triazol-4-yl)benzyl)-3-methylurea (6)

After Boc deprotection of **27** using general procedure B, the free amine (0.182 mmol) was stirred with triethylamine (43.7 µL, 0.313 mmol) in dichloromethane (1.25 mL) and stirred for 30 minutes. In a separate vessel, triphosgene (18.6 mg, 0.063 mmol) was dissolved in dichloromethane (1.25 mL) and cooled to 0 °C. The amine solution was added dropwise to the triphosgene solution and the mixture was warmed to room temperature and stirred for 30 minutes. The solution was cooled to 0 °C and 1.25 M methylamine in methanol (156 µL, 1.25 mmol) was added and stirred for 1 hour. When the reaction was complete by LC/MS analysis, the solvent was removed *in vacuo* and the product purified using reverse phase column chromatography. Yield: 11.8 mg, 25%, white solid. ^1^H NMR (399 MHz, CD_3_OD) δ 7.66 – 7.57 (m, 2H), 7.55 – 7.47 (m, 1H), 7.28 (d, *J* = 7.7 Hz, 1H), 3.97 (s, 2H), 2.67 (s, 3H), 2.21 (s, 6H). ^13^C NMR (100 MHz, CD_3_OD) δ 153.5, 139.2, 132.7, 132.0, 130.5, 130.0, 129.3, 111.4, 40.5, 27.0, 10.6. IR (neat) *ṽ* (cm^-1^): 3260, 1690, 1545, 1495, 1416, 1242, 786. HRMS (DART) *m/z*: [M + H]^+^ calcd for C_13_H_18_N_5_O 260.1511, found 260.1510.

### *N*-(2-(3,5-dimethyl-4*H*-1,2,4-triazol-4-yl)benzyl)methanesulfonamide (7)

After Boc deprotection of **27** using general procedure B, free amine (1.0 equiv., 48 mg, 0.237 mmol) and triethylamine (1.2 equiv., 39.7 μL, 0.285 mmol) were dissolved in anhydrous dichloromethane (2.4 mL, 0.1 M) in an oven-dried 2-dram vial equipped with magnetic stir bar and rubber septum. The resulting mixture was cooled to 0 °C and then methanesulfonyl chloride (1.2 equiv., 22 μL, 0.285 mmol) was added. After stirring at 0 °C for 1 hour, the reaction was warmed to room temperature over 2 hours and then quenched with 5 mL of a saturated solution of sodium bicarbonate. The organic layer was separated and the aqueous layer was extracted with dichloromethane (3×5 mL). The organic extracts were combined, dried over sodium sulfate, filtered, and solvent was removed *in vacuo*. The crude residue was then purified by reverse-phase flash-chromatography using a 5-100% MeCN gradient in H_2_O with 1% of formic acid. The fractions containing desired product were pooled and lyophilized. Yield: 35%, white solid. ^1^H NMR (400 MHz, CD_3_OD) δ 2.10 - 2.30 (m, 6H), 2.86 (s, 3H), 3.95 (s, 2H), 7.32 (d, *J* = 7.7 Hz, 1H), 7.54 - 7.62 (m, 1H), 7.62 - 7.69 (m, 1H), 7.75 (d, *J* = 7.7 Hz, 1H), 8.46 (s, sulfonamide N**H** under deuterium exchange). ^13^C NMR (126 MHz, CD_3_OD) δ 153.6, 137.1, 133.2, 132.2, 132.0, 130.8, 129.6, 43.4, 40.0, 10.8; HRMS (DART) m/z: [M + H]+ calcd for C_12_H_17_N_4_O_2_S 281.1072, found 281.1070.

### 2-(3,5-dimethyl-4*H*-1,2,4-triazol-4-yl)benzoic acid (8)

Anhydride **28** ^21^ (1.0 g, 6.21 mmol, 1.1 equiv.) and acetylhydrazide (418 mg, 5.64 mmol, 1.0 equiv.) were refluxed in EtOH (56 mL, 0.1M) for 3 h.^22^ The reaction was cooled down to r.t. and then dry loaded on celite. The crude was purified using C18 functionlized silica gel using a gradient from 5:95 MeCN:H_2_O to 1:1 MeCN:H_2_O. The fractions containing the desired product were lyophilized. Yield: 27%, white solid. ^1^H NMR (500 MHz, DMSO-*d*_6_) δ 13.38 (s, 1H), 8.05 (dd, *J* = 7.8, 1.6 Hz, 1H), 7.78 (dt, *J* = 7.7, 1.7 Hz, 1H), 7.68 (dt, *J* = 7.6, 1.3 Hz, 1H), 7.49 (dd, *J* = 7.8, 1.3 Hz, 1H), 2.01 (s, 6H). ^13^C NMR (126 MHz, DMSO-*d_6_*) δ 165.8, 150.5, 133.6, 133.0, 131.5, 130.3, 129.6, 129.6, 10.6. HRMS (DART) *m/z*: [M + H]^+^ calcd for C_11_H_12_N_3_O_2_ 218.0930, found 218.0930.

### *tert*-butyl (2-acetamido-5-bromobenzyl)carbamate (29a)

Synthesized according to general procedure D. Scale: 1.89 mmol. Yield: 569 mg, 88%, white solid. *R_f_* (1:3 EtOAc:hexanes): 0.42. ^1^H NMR (500 MHz, CDCl_3_) δ 9.51 (br. s, 1H), 8.17 (d, *J* = 8.8 Hz, 1H), 7.40 (dd, *J* = 8.8, 2.4 Hz, 1H), 7.29 – 7.26 (m, 1H), 5.09 (br. s, 1H), 4.22 (d, *J* = 6.9 Hz, 2H), 2.22 (s, 3H), 1.45 (s, 9H). ^13^C NMR (126 MHz, CDCl_3_) δ 169.5, 157.5, 136.4, 133.1, 132.0, 130.2, 124.0, 116.2, 81.2, 41.5, 28.5, 24.6. IR (neat) *ṽ* (cm^−1^): 3360, 3251, 2988, 1686, 1666, 1539, 1514, 1278, 1270, 1163, 641. HRMS (DART) *m/z*: [M + H]^+^ calcd for C_14_H_20_BrN_2_O_3_ 343.0665, found 343.0664.

### *tert*-butyl (2-acetamido-5-chlorobenzyl)carbamate (29b)

Synthesized according to procedure D. Scale: 1.89 mmol. Yield: 295 mg, 52%, white solid. *R_f_* (1:2 EtOAc:hexanes): 0.35. ^1^H NMR (500 MHz, CDCl_3_) δ 9.50 (s, 1H), 8.21 (d, *J* = 8.8 Hz, 1H), 7.26 (dd, *J* = 8.8, 2.5 Hz, 1H), 7.11 (d, *J* = 2.5 Hz, 1H), 5.10 (s, 1H), 4.22 (d, *J* = 6.8 Hz, 2H), 2.23 (s, 3H), 1.45 (s, 9H). ^13^C NMR (126 MHz, CDCl_3_) *δ* 171.2, 157.5, 135.8, 130.2, 130.0, 129.0, 128.6, 123.7, 81.2, 77.2, 41.6, 28.5, 24.5. IR (neat) *ṽ* (cm^-1^): 3357, 3254, 2982, 1665, 1542, 1515, 1365, 1277, 1162, 932, 837. HRMS (DART) *m/z*: [M + H]^+^ calcd for C_14_H_20_ClN_2_O_3_ 298.1084, found 298.1082.

### *tert*-butyl (5-bromo-2-ethanethioamidobenzyl)carbamate (30a)

Synthesized according to general procedure A. Scale: 23.3 mmol. Yield: 6.93 g, 83%, pale yellow powder. *R_f_* (1:3 EtOAc:hexanes): 0.5. ^1^H NMR (500 MHz, CDCl_3_) δ 11.24 (br. s, 1H), 8.08 (d, *J* = 8.7 Hz, 1H), 7.46 (dd, *J* = 8.7, 2.3, Hz, 1H), 7.39 (d, *J* = 2.3 Hz, 1H), 5.25 (br. t, *J* = 5.6 Hz, 1H), 4.12 (d, *J* = 6.7 Hz, 2H), 2.79 (d, *J* = 0.5 Hz, 3H), 1.42 (s, 9H). ^13^C NMR (126 MHz, CDCl_3_) δ 201.8, 157.6, 136.8, 134.3, 133.4, 131.3, 128.4, 120.0, 81.4, 40.7, 35.4, 28.4. IR (neat) *ṽ* (cm^-1^): 3257, 2934, 1664, 1530, 1374, 1287, 1157, 701. HRMS (DART) *m/z*: [M + H]^+^ calcd for C_14_H_20_BrN_2_O_2_S 359.0429, found 359.0431.

### *tert*-butyl (5-chloro-2-ethanethioamidobenzyl)carbamate (30b)

Synthesized according to general procedure A. Scale: 0.94 mmol. Yield: 174 mg, 60%, pale yellow solid. *R_f_* (1:3 EtOAc:hexanes): 0.5. ^1^H NMR (500 MHz, CDCl_3_) δ 11.22 (br. s, 1H), 8.13 (br. d, *J* = 8.7 Hz, 1H), 7.32 (ddd, *J* = 8.7, 2.4, 0.4 Hz, 1H), 7.24 (d, *J* = 2.4 Hz, 1H), 5.23 (br. t, *J* = 6.7 Hz, 1H), 4.13 (d, *J* = 6.7 Hz, 2H), 2.80 (d, *J* = 0.5 Hz, 3H), 1.43 (s, 9H). ^13^C NMR (126 MHz, CDCl_3_) δ 201.8, 157.6, 136.3, 134.0, 132.1, 130.4, 128.4, 128.2, 81.4, 77.2, 40.8, 35.4, 28.4. IR (neat) *ṽ* (cm^-1^): 3267, 2934, 1669, 1530, 1381, 1366, 1288, 1154, 854, 746, 701. HRMS (DART) *m/z:* [M + H]^+^ calcd for C_14_H_20_ClN_2_O_2_S 315.0934, found 315.0927.

### *tert*-butyl (5-bromo-2-(3,5-dimethyl-4*H*-1,2,4-triazol-4-yl)benzyl)carbamate (31a)

Synthesized according to general procedure C. Scale: 0.56 mmol. Yield: 190.2 mg, 90%, white solid. *R_f_* (1:19 MeOH:EtOAc): 0.4. ^1^H NMR (500 MHz, CDCl_3_) δ 7.73 (d, *J* = 2.2 Hz, 1H), 7.58 (dd, *J* = 8.2, 2.2 Hz, 1H), 6.98 (d, *J* = 8.2 Hz, 1H), 5.07 (br. s, 1H), 3.91 (d, *J* = 6.2 Hz, 2H), 2.20 (s, 6H), 1.42 (s, 9H). ^13^C NMR (126 MHz, CDCl_3_) δ 155.7, 151.2, 139.1, 132.9, 132.3, 131.3, 129.6, 125.0, 80.5, 40.4, 28.4, 11.1. IR (neat) *ṽ* (cm^-1^): 3251, 2976, 1706, 1529, 1270, 1249, 1161, 874, 826. HRMS (DART) *m/z*: [M + H]^+^ calcd for C_16_H_22_BrN_4_O_2_ 381.0926, found 381.0928.

### *tert*-butyl (5-chloro-2-(3,5-dimethyl-4*H*-1,2,4-triazol-4-yl)benzyl)carbamate (31b)

Synthesized according to general procedure C. Scale: 0.51 mmol. Yield: 95.7 mg, 56%, white solid. *R_f_* (1:19 MeOH:EtOAc): 0.41. ^1^H NMR (500 MHz, CDCl_3_) δ 7.58 (d, *J* = 2.3 Hz, 1H), 7.42 (dd, *J* = 8.2, 2.3 Hz, 1H), 7.04 (d, *J* = 8.2 Hz, 1H), 5.18 (br. s, 1H), 3.93 (d, *J* = 6.2 Hz, 2H), 2.20 (s, 6H), 1.42 (s, 9H). ^13^C NMR (126 MHz, CDCl_3_) δ 155.8, 151.5, 138.9, 136.8, 130.8, 130.0, 129.4, 129.3, 80.4, 40.6, 28.5, 11.1. IR (neat) *ṽ* (cm^-1^): 3250, 2972, 1705, 1532, 1269, 1250, 1162, 854, 747. HRMS (DART) *m/z*: [M + H]^+^ calcd for C_16_H_22_ClN_4_O_2_ 337.1423, found 337.1421.

### *N*-(5-bromo-2-(3,5-dimethyl-4*H*-1,2,4-triazol-4-yl)benzyl)acetamide (9a)

Synthesized according to general procedure B. Scale: 0.41 mmol. Yield: 101.3 mg, 77%, white solid. *R_f_* (1:9 MeOH:CHCl_3_): 0.2. ^1^H NMR (500 MHz, CDCl_3_) δ 7.87 (d, *J* = 2.3 Hz, 1H), 7.56 (dd, *J* = 8.3, 2.3 Hz, 1H), 7.47 (br. s, 1H), 6.94 (d, *J* = 8.3 Hz, 1H), 4.11 (d, *J* = 6.0 Hz, 2H), 2.18 (s, 6H), 2.02 (s, 3H). ^13^C NMR (126 MHz, CDCl_3_) δ 170.9, 151.5, 138.8, 134.5, 132.4, 131.4, 129.5, 124.9, 39.7, 23.0, 11.1. IR (neat) *ṽ* (cm^-1^): 3257, 3043, 2931, 1645, 1532, 1496, 1417, 1372, 1023, 1009, 868, 629. HRMS (DART) *m/z*: [M + H]^+^ calcd for C_13_H_16_BrN_4_O 323.0508, found 323.0513.

### *N*-(5-chloro-2-(3,5-dimethyl-4*H*-1,2,4-triazol-4-yl)benzyl)acetamide (9b)

Synthesized according to general procedure B. Scale: 0.26 mmol. Yield: 61.6 mg, 87%, white solid. *R_f_* (1:9 MeOH:CHCl_3_): 0.2. ^1^H NMR (500 MHz, CDCl_3_) δ 7.74 (d, *J* = 2.4 Hz, 1H), 7.70 (br. s, 1H), 7.40 (dd, *J* = 8.3, 2.4 Hz, 1H), 7.00 (d, *J* = 8.3 Hz, 1H), 4.13 (d, *J* = 5.9 Hz, 2H), 2.17 (s, 6H), 2.03 (s, 3H). ^13^C NMR (126 MHz, CDCl_3_) δ 171.0, 151.5, 138.7, 136.8, 131.8, 130.9, 129.4, 129.2, 39.9, 23.0, 11.1. IR (neat) *ṽ* (cm^-1^): 3248, 3046, 1659, 1646, 1534, 1498, 1419, 1373, 1024, 870. HRMS (DART) *m/z*: [M + H]^+^ calcd for C_13_H_16_ClN_4_O 279.1013, found 279.1015.

### *N*-(5-cyano-2-(3,5-dimethyl-4*H*-1,2,4-triazol-4-yl)benzyl)acetamide (9c)

Bromide **9a** (20 mg, 0.062 mmol), sodium carbonate (6.56 mg, 0.062 mmol), palladium acetate (0.695 mg, 3.09 µmol), and isopropanol (4.8 µL), were added to degassed DMF (100 µL) and heated to 140 °C. Ground up K_4_[Fe(CN)_6_]•3H_2_O (7.84 mg, 0.019 mmol) was added, the vessel was sealed and heated at 140 °C overnight. The solvent was evaporated *in vacuo* and the crude product purified by preparatory TLC. Yield: 6.0 mg, 36%, white solid. *R_f_* (3:17 MeOH:CHCl_3_): 0.35. ^1^H NMR (500 MHz, CDCl_3_) δ 8.08 (d, *J* = 1.9 Hz, 1H), 7.75 (dd, *J* = 8.0, 1.9 Hz, 1H), 7.68 (s, 1H), 7.22 (d, *J* = 8.0 Hz, 1H), 4.18 (d, *J* = 5.8 Hz, 2H), 2.20 (s, 6H), 2.04 (s, 3H). ^13^C NMR (126 MHz, CDCl_3_) δ 171.1, 151.2, 138.8, 136.2, 135.5, 132.7, 129.1, 117.4, 115.1, 39.8, 22.9, 11.1. IR (neat) *ṽ* (cm^-1^): 3254, 3037, 2928, 2235, 1677, 1658, 1534, 1501, 1294, 1013, 847. HRMS (DART) *m/z*: [M + H]^+^ calcd for C_14_H_16_N_5_O 270.1355, found 270.1350.

### *N*-(2-(3,5-dimethyl-4*H*-1,2,4-triazol-4-yl)-5-methylbenzyl)acetamide (9d)

Bromide **9a** (10.0 mg, 0.031 mmol), potassium carbonate (12.8 mg, 0.093 mmol), methyl boronic acid (5.6 mg, 0.093 mmol), and Pd(PPh_3_)_4_ (3.6 mg, 3.09 µmol) were added to a vial under N_2_ and degassed 9:1 dioxane:water (154.5 µL) was added. The vessel was sealed and heated to 100 °C for 20 hours. The reaction was diluted with chloroform and washed with water and brine and solvent was removed *in vacuo* and the crude product was purified by column chromatography. Yield: 5.6 mg, 70%, white solid. *R_f_* (1:9 MeOH:CHCl_3_): 0.1. ^1^H NMR (500 MHz, CDCl_3_) δ 7.66 (br. t, *J* = 6.0 Hz, 1H), 7.51 (d, *J* = 1.9 Hz, 1H), 7.23 – 7.17 (m, 1H), 6.91 (d, *J* = 7.9 Hz, 1H), 4.11 (d, *J* = 6.0 Hz, 2H), 2.42 (s, 3H), 2.15 (s, 6H), 2.00 (s, 3H). ^13^C NMR (126 MHz, CDCl_3_) δ 170.9, 151.7, 141.0, 136.2, 132.2, 129.8, 129.7, 127.6, 39.9, 23.0, 21.5, 11.1. IR (neat) *ṽ* (cm^-1^): 3212, 3070, 2967, 1676, 1537, 1504, 1421, 1025, 829. HRMS (DART) *m/z*: [M + H]^+^ calcd for C_14_H_19_N_4_O 259.1559, found 259.1560.

### *N*-(5-cyclopropyl-2-(3,5-dimethyl-4*H*-1,2,4-triazol-4-yl)benzyl)acetamide (9e)

Bromide **9a** (15 mg, 0.046 mmol), potassium carbonate (19.2 mg, 0.14 mmol), cyclopropyl boronic acid (12.0 mg, 0.14 mmol), and Pd(PPh_3_)_4_ (5.36 mg, 4.64 µmol) were added to a vial under N_2_ and degassed 9:1 dioxane:water (232 µL) was added. The vessel was sealed and heated to 100 °C for 20 hours. The reaction was diluted with chloroform and washed with water and brine and solvent was removed *in vacuo* and the crude product was purified by column chromatography. Yield: 7.7 mg, 58%, white solid. *R_f_* (1:9 MeOH:CHCl_3_): 0.1. ^1^H NMR (500 MHz, CDCl_3_) δ 7.39 (d, *J* = 2.1 Hz, 1H), 7.05 (dd, *J* = 8.1, 2.1 Hz, 1H), 7.02 (br. t, *J* = 6.0 Hz, 1H), 6.93 (d, *J* = 8.1 Hz, 1H), 4.10 (d, *J* = 6.0 Hz, 2H), 2.18 (s, 6H), 2.00 (s, 3H), 1.96 (tt, *J* = 8.4, 5.3 Hz, 1H), 1.09 – 1.04 (m, 2H), 0.80 – 0.76 (m, 2H). ^13^C NMR (126 MHz, CDCl_3_) δ 170.7, 151.9, 147.4, 136.1, 129.4, 128.8, 127.8, 125.8, 39.9, 23.1, 15.5, 11.2, 10.3. IR (neat) *ṽ* (cm^-1^): 3236, 3067, 2922, 1638, 1560, 1536, 1511, 1418, 1375, 1291, 1015. HRMS (DART) *m/z*: [M + H]^+^ calcd for C_16_H_21_N_4_O 285.1715, found 285.1716.

### 2-(3-(((*tert*-butoxycarbonyl)amino)methyl)-5-methyl-4*H*-1,2,4-triazol-4-yl)benzoic acid (33)

Anhydride **28** (2.65 g, 16.4 mmol, 1.0 equiv.)^21^ and hydrazide **32** (mg, mmol, 1.2 equiv.) ^22^ were refluxed EtOH for 3 hours.^23^ The reaction was cooled down to room temperature and the reaction was dry loaded in celite and purified on C18 functionalized silica column using a gradient from 5:95 MeCN:H_2_O to 1:1 MeCN:H_2_O. The fractions containing the product were combined and lyophilized to give the product. Yield: 3.1g, 57%, white solid. ^1^H NMR (500 MHz, DMSO-*d*_6_) δ 13.30 (s, 1H), 8.07 (dd, J = 7.8, 1.6 Hz, 1H), 7.81 (dt, J = 7.6, 1.7 Hz, 1H), 7.71 (dt, J = 7.6, 1.3 Hz, 1H), 7.54 (dd, J = 7.8, 1.3 Hz, 1H), 6.74 (t, J = 5.9 Hz, 1H), 3.18 – 3.06 (m, 2H), 2.02 (s, 3H), 1.32 (s, 9H). ^13^C NMR (126 MHz, DMSO-*d*_6_) δ 165.5, 154.8, 151.8, 151.3, 133.5, 132.8, 131.6, 130.2, 129.7, 128.9, 77.9, 35.0, 28.1, 10.4. HRMS (DART) *m/z*: [M + H]^+^ calcd for C_16_H_21_N_4_O4 333.1563, found 333.1565.

### *tert*-butyl ((4-(2-(hydroxymethyl)phenyl)-5-methyl-4*H*-1,2,4-triazol-3- yl)methyl)carbamate (34)

Compound **33** (2.7 g, 8.12 mmol, 1.0 equiv.) was dissolved in freshly distilled THF (81 mL, 0.1 M) and the reaction kept under a nitrogen atmosphere. LAH (1.21 g, 31.9 mmol, 3.9 equiv.) was then added in one portion and the resulting mixture was heated to 40°C overnight. The next day the reaction was cooled down to room temperature and the LAH was quenched using a Fieser workup. After filtration, the resulting filtrate was dry loaded into celite and purified using C18 functionalized silica using a gradient from 5:95 MeCN:H_2_O to 95:5 MeCN:H_2_O. The fractions containing the product were combined and lyophilized to give the desired product. Yield: 1.4 g, 54%, white solid. ^1^H NMR (399 MHz, DMSO-*d*_6_) δ 7.63 (dd, *J* = 7.8, 1.6 Hz, 1H), 7.58 – 7.51 (m, 1H), 7.46 (dt, *J* = 7.6, 1.7 Hz, 1H), 7.28 (dd, *J* = 7.7, 1.3 Hz, 1H), 7.14 (t, *J* = 5.9 Hz, 1H), 5.22 (t, *J* = 5.3 Hz, 1H), 4.18 – 4.05 (m, 2H), 4.05 – 3.91 (m, 2H), 2.05 (s, 3H), 1.28 (s, 9H). ^13^C NMR (126 MHz, DMSO-*d*_6_) δ 155.4, 152.4, 152.0, 139.4, 131.6, 130.6, 129.7, 129.0, 128.4, 78.5, 59.3, 35.4, 28.5, 10.7. HRMS (DART) *m/z*: [M + H]^+^ calcd for C_16_H_21_N_4_O_4_ 319.1770, found 319.1769.

### *tert*-butyl ((4-(2-((1,3-dioxoisoindolin-2-yl)methyl)phenyl)-5-methyl-4*H*-1,2,4-triazol-3- yl)methyl)carbamate (35)

Compound **34** (1.01g, 3.17 mmol, 1.0 equiv.), phthalamide (583 mg, 3.97 mmol, 1.25 equiv.), and triphenylphosphine (1.04g, 3.97 mmol, 1.25 equiv.) were dissolved in freshly distilled THF (32 mL, 0.1 M) and the mixture kept under a nitrogen atmosphere. DIAD was added in one portion and stirred overnight at r.t. under a nitrogen atmosphere. The mixture was dry loaded in silica gel and purified in silica gel eluting a solution of 2.5% MeOH in dichloromethane slowly increasing the gradient until 5% MeOH in DCM was reached. Yield: 1.36g, 96%, white solid. ^1^H NMR (500 MHz, DMSO-*d*_6_) δ 7.90 – 7.81 (m, 4H), 7.48 – 7.44 (m, 2H), 7.37 (dd, *J* = 6.7, 2.7 Hz, 1H), 7.34 – 7.30 (m, 1H), 7.20 (t, *J* = 5.7 Hz, 1H), 4.59 (d, *J* = 16.1 Hz, 1H), 4.24 – 4.12 (m, 2H), 4.11 – 4.02 (m, 1H), 2.17 (s, 3H), 1.27 (s, 9H). ^13^C NMR (126 MHz, DMSO-*d*_6_) δ 168.1, 155.3, 152.6, 152.1, 135.1, 134.4, 131.9, 131.8, 130.9, 129.4, 129.0, 128.9, 123.7, 78.5, 36.8, 35.4, 28.5, 10.7. HRMS (DART) *m/z*: [M + H]^+^ calcd for C_24_H_26_N_5_O_4_ 448.1985, found 448.1988.

### *tert*-butyl ((4-(2-(acetamidomethyl)phenyl)-5-methyl-4*H*-1,2,4-triazol-3- yl)methyl)carbamate (36)

Compound **35** (1.2 g, 2.68 mmol, 1.0 equiv.) and hydrazine monohydrate (523 μL, 10.7 mmol, 4.0 equiv.) were refluxed for 6 hours in MeOH (27 mL, 0.1 M). The reaction was monitored by LC-MS. Once the reaction was completed, the reaction was cooled to r.t. and then *N*-acetyl-*N*-methoxyacetamide (2.11 g, 16.1 mmol, 6.0 equiv.) were added to the reaction mixture and stirred overnight at room temperature. The resulting reaction mixture was dry loaded in celite and purified on C18 functionalized silica column using a gradient from 5:95 MeCN:H_2_O to 95:5 MeCN:H_2_O. Yield: 490 mg, 51% white solid. ^1^H NMR (500 MHz, DMSO- *d*_6_) δ 8.37 (t, *J* = 5.9 Hz, 1H), 7.55 – 7.45 (m, 2H), 7.41 (dt, *J* = 7.5, 1.9 Hz, 1H), 7.26 (dd, *J* = 7.8, 1.3 Hz, 1H), 7.14 (t, *J* = 5.8 Hz, 1H), 4.10 – 3.93 (m, 2H), 3.83 (d, *J* = 5.9 Hz, 2H), 2.07 (s, 3H), 1.81 (s, 3H), 1.26 (s, 9H). ^13^C NMR (126 MHz, DMSO-*d*_6_) δ 169.3, 155.0, 151.9, 151.5, 136.9, 131.3, 130.1, 129.0, 128.3, 128.1, 78.0, 37.5, 35.0, 28.1, 22.4, 10.3. HRMS (DART) *m/z*: [M + H]^+^ calcd for C_18_H_25_N_5_O_3_ 360.2036, found 360.2038.

### *N*-(2-(3-(aminomethyl)-5-methyl-4*H*-1,2,4-triazol-4-yl)benzyl)acetamide (37)

Compound 36 (251 mg, 0.70 mmol, 1.0 equiv.) was stirred for 45 min in a 1:1 DCM:TFA mixture. Nitrogen was blown into the reaction to remove solvents. The resulting crude was redissolved and neutralized with IRA-67 resin and then dry loaded in celite and purified using C18 functionalized silica using a gradient from 0:95 MeCN:H_2_O to 1:4 MeCN:H_2_O. Yield: 121 mg, 67% white solid.^1^H NMR (500 MHz, CD_3_OD) δ 7.66 – 7.58 (m, 2H), 7.53 (ddd, *J* = 7.7, 7.1, 2.0 Hz, 1H), 7.35 (dd, *J* = 7.8, 1.2 Hz, 1H), 4.15 – 3.95 (m, 2H), 3.68 (s, 2H), 2.23 (s, 3H), 1.93 (s, 3H). ^13^C NMR (126 MHz, CD_3_OD) δ 173.0, 157.0, 154.0, 137.8, 132.4, 132.2, 130.9, 130.3, 129.5, 39.6, 36.9, 22.3, 10.5. HRMS (DART) *m/z*: [M + H]^+^ calcd for C_13_H_18_N_5_O 260.1512, found 260.1511.

### *N*-(*N*-(*N*^2^-(*tert*-butoxycarbonyl)-*N*^6^,*N*^6^-dimethyl-*L*-lysyl)-*O*-(*tert*-butyl)-*L*-seryl)-*O*-(*tert*- butyl)-*L*-threonylglycine (40)

Tetrapeptide **38** was synthesized using standard chloro-trityl resin for solid phase peptide synthesis using 1.0 g of resin (1 mmol). **38** (400 mg, 0.542 mmol, 1.0 equiv.) was dissolved in 1.1 mL of MeOH (0.5 M) and added AcOH (124 μL, 2.17 mmol, 4.0 equiv.) and the solution was charged in Pd on carbon (29 mg, 0.027 mmol, 5 mol%). The resulting solution was stirred under a H_2_ atmosphere for 12 h. The reaction was monitored by LC-MS. Upon completion of the reaction, the resulting crude was filtered through celite and the product concentrated to give peptide **39** in quantitative yield. Peptide **39** was then dissolved in MeOH and formaldeyde (149 μL, 1.63 mmol, 3.0 equiv., 37% aq. solution) was added and stirred 20 min at room temperature. NaBH_3_CN (136 mg, 2.17 mmol, 4.0 equiv.) was added and the reaction stirred for 8 h at room temperature. The reaction was slowly quenched with 1 M HCl until no more bubbling was observed. The resulting mixture was then dry loaded in celite and purified using C18 functionalized silica column using a gradient from 5:95 MeCN:H_2_O to 95:5 MeCN:H_2_O with 0.1% formic acid as an additive. The fractions containing **40** were lyophilized. Yield: 291 mg, 85%, white solid. ^1^H NMR (500 MHz, DMSO-*d*_6_) δ 8.17 (d, *J* = 8.0 Hz, 1H), 7.61 (t, *J* = 5.5 Hz, 1H), 7.37 (d, *J* = 7.7 Hz, 1H), 6.87 (d, *J* = 8.2 Hz, 1H), 4.35 (dt, *J* = 8.7, 4.6 Hz, 1H), 4.15 (dd, *J* = 7.7, 3.2 Hz, 1H), 4.05 (dt, *J* = 8.3, 4.8 Hz, 1H), 3.99 (dq, *J* = 6.3, 3.1 Hz, 1H), 3.68 – 3.54 (m, 2H), 3.54 – 3.38 (m, 2H), 2.75 (dt, *J* = 12.0, 7.7 Hz, 1H), 2.59 – 2.53 (m, 2H), 2.47 (s, 6H), 1.64 (dd, *J* = 15.2, 5.9 Hz, 1H), 1.58 – 1.45 (m, 3H), 1.37 (s, 9H), 1.27 (ddd, *J* = 21.3, 11.4, 5.8 Hz, 1H), 1.13 (d, *J* = 1.7 Hz, 19H), 0.99 (d, *J* = 6.3 Hz, 3H). ^13^C NMR (126 MHz, DMSO-*d*_6_) δ 172.7, 172.5, 170.0, 169.1, 164.1, 155.7, 78.4, 74.2, 73.4, 66.8, 62.1, 58.1, 57.2, 54.1, 54.1, 43.2, 42.9, 40.9, 31.8, 28.6, 28.3, 27.5, 24.7, 23.0, 19.2. HRMS (DART) *m/z*: [M + H]^+^ calcd for C_35_H_58_N_5_O_9_ 632.4235, found 632.4242.

### (*S*)-*N*-((*S*)-1-(((2*S*,3*R*)-1-((2-(((4-(2-(acetamidomethyl)phenyl)-5-methyl-4*H*-1,2,4-triazol-3-yl)methyl)amino)-2-oxoethyl)amino)-3-hydroxy-1-oxobutan-2-yl)amino)-3-hydroxy-1- oxopropan-2-yl)-2-amino-6-(dimethylamino)hexanamide (10)

Compound **40** (155 mg, 0.245 mmol, 1.0 equiv.) was dissolved in 2.5 mL of DCM (0.1M) and EDC•HCl (70 mg, 0.368 mmol, 1.5 equiv.) was added in one portion and stirred the resulting mixture for 20 min. Amine **37** (95 mg, 0.368 mmol, 1.5 equiv.) was added and the reaction stirred for 6 h at room temperature. The progress of the reaction was monitored by LC-MS. Upon complete consumption of **40**, the reaction was dry loaded in celite and quickly purified using C18 functionalized silica using a gradient from 5:95 MeCN:H_2_O to 95:5 MeCN:H_2_O with 0.1% formic acid as an additive. The fractions containing coupled product were lyophilized and then treated with 2.5 mL a 95:2.5:2.5 TFA:H_2_O:triethylsilane for 3 h. TFA was then removed by blowing nitrogen into the reaction vessel. The resulting concentrate was redissolved in MeOH and IRA-67 resin was added until the pH of the solution reached ~ 8. This mixture containing the resin was then dry loaded in celite dry and purified using C18 functionalized silica using a gradient from 0:100 MeCN:H_2_O to 15:85 MeCN:H_2_O (no formic acid as additive in order to isolate the free amine). The fractions containing **10** were lyophilized for isolation. Yield: 48 mg, 30%, white solid. ^1^H NMR (500 MHz, DMSO- *d*_6_) δ 8.67 (t, *J* = 7.2 Hz, 1H), 8.45 (dt, *J* = 5.9, 1.4 Hz, 1H), 8.30 – 8.23 (m, 2H), 8.00 (dt, *J* = 7.6, 5.8 Hz, 1H), 7.86 (dd, *J* = 8.1, 3.0 Hz, 1H), 7.55 (dt, *J* = 7.5, 1.3 Hz, 1H), 7.50 – 7.47 (m, 1H), 7.44 (dt, *J* = 7.6, 1.6 Hz, 1H), 7.30 (ddd, *J* = 7.8, 3.2, 1.3 Hz, 1H), 4.46 (q, *J* = 6.1 Hz, 1H), 4.20 (ddd, *J* = 8.0, 6.0, 3.5 Hz, 1H), 4.17 – 4.09 (m, 2H), 4.05 (ddd, *J* = 6.2, 3.6, 2.4 Hz, 1H), 3.90 – 3.78 (m, *J* = 6.0 Hz, 2H), 3.77 – 3.64 (m, 2H), 3.64 – 3.55 (m, 3H), 2.60 (dt, *J* = 7.2, 2.4 Hz, 2H), 2.43 (s, 6H), 2.11 (s, 3H), 1.83 (d, *J* = 2.4 Hz, 3H), 1.67 (ddq, *J* = 36.5, 14.0, 7.1 Hz, 2H), 1.50 (ddd, *J* = 13.4, 8.0, 4.4 Hz, 2H), 1.34 (p, *J* = 7.3, 6.8 Hz, 2H), 1.03 (dd, *J* = 6.4, 1.5 Hz, 3H). ^13^C NMR (126 MHz, DMSO-*d*_6_) δ 170.34, 170.3, 170.2, 170.2, 170.0, 169.4, 169.4, 168.4, 151.6, 151.3, 136.8, 136.8, 131.5, 131.5, 131.0, 131.0, 130.3, 128.8, 128.8, 128.4, 128.2, 66.3, 61.6, 58.4, 58.3, 57.2, 55.0, 55.0, 52.5, 43.3, 41.8, 37.6, 33.6, 31.7, 24.8, 22.4, 21.8, 19.7, 19.7, 10.3. HRMS (DART) *m/z*: [M + H]^+^ calcd for C_30_H_49_N_10_O_7_ 661.3780, found 661.3772.

## Acknowledgements

This research was supported by the Natural Sciences and Engineering Research Council of Canada (2015-05939 to CHA, and Collaborative Research and Training Experience 432008-2013). The SGC is a registered charity (number 1097737) that receives funds from AbbVie, Bayer Pharma AG, Boehringer Ingelheim, Canada Foundation for Innovation, Eshelman Institute for Innovation, Genome Canada through Ontario Genomics Institute, Innovative Medicines Initiative (EU/EFPIA) [ULTRA-DD grant no. 115766], Janssen, Merck & Co., Novartis Pharma AG, Ontario Ministry of Economic Development and Innovation, Pfizer, São Paulo Research Foundation-FAPESP, Takeda, and the Wellcome Trust.

---

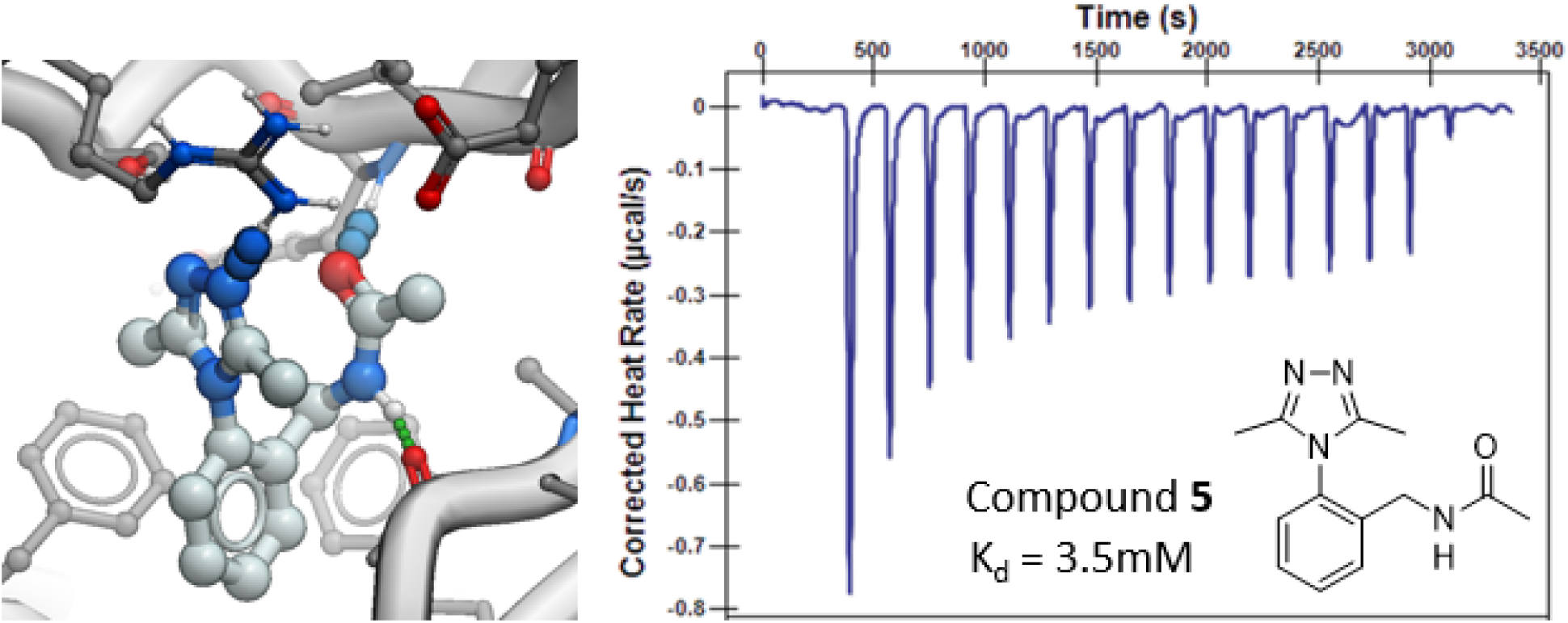
TABLE OF CONTENTS GRAPHIC

