## Supplemental Data for "Identification and Characterization of the First Fragment Hits for SETDB1 Tudor Domain"

**Supplementary Figure 1**: Constrained analogues of known bromodomain inhibitors tested by crystal soaking

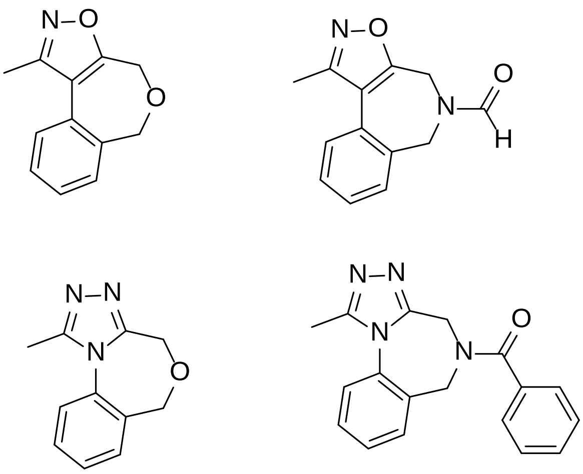

**Supplementary Figure 2**: HSQC NMR of SETDB1 TTD with fragments and DMSO (control)

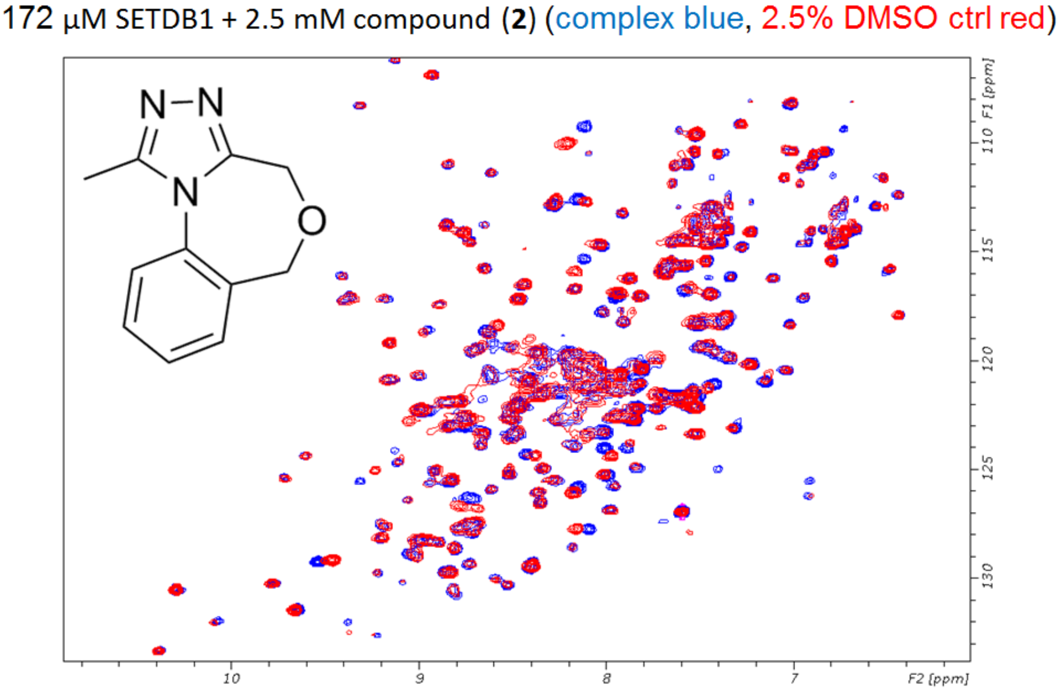

200 µM SETDB1 + 10 mM compound (**4**) (complex blue, 2.5% DMSO ctrl red)

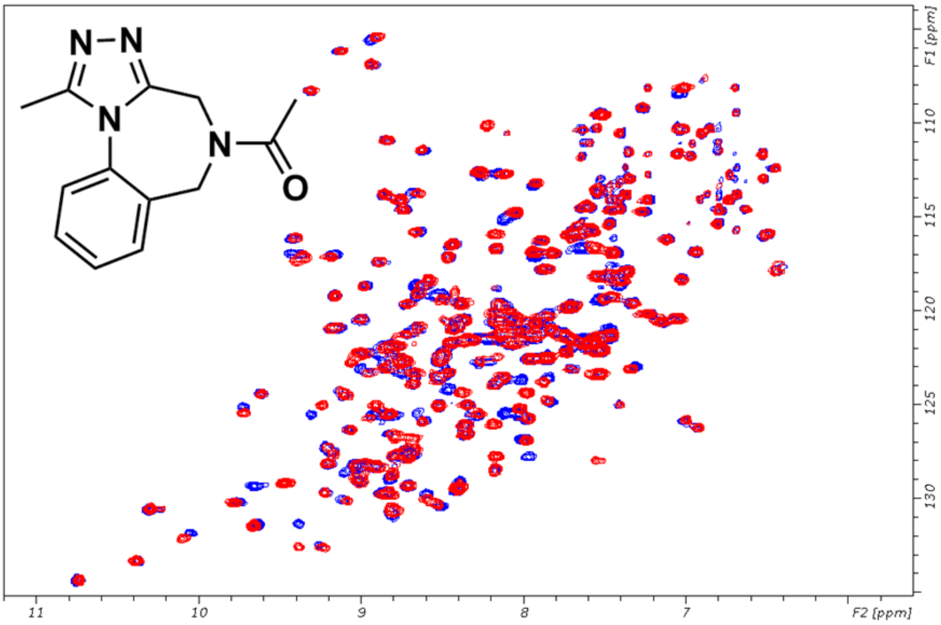


200 µM SETDB1 + 10 mM compound (**5**) (complex blue, 2.5% DMSO ctrl red)

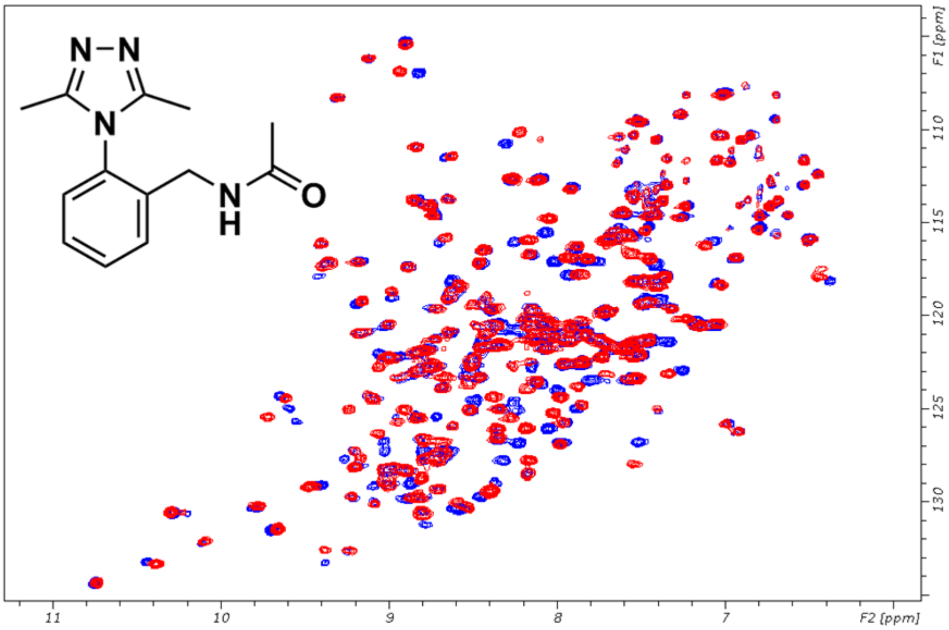

135 μM SETDB1 + 5 mM compound (**7**) (complex blue, 2.5% DMSO ctrl red)

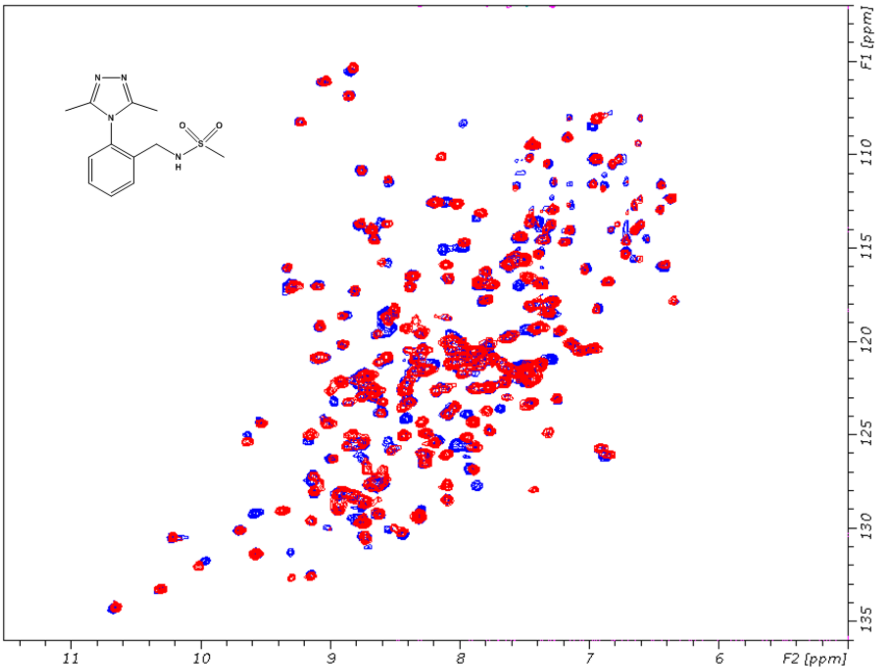


200 μM SETDB1 + 5 mM compound (**10**) (complex blue, 2.5% DMSO ctrl red)

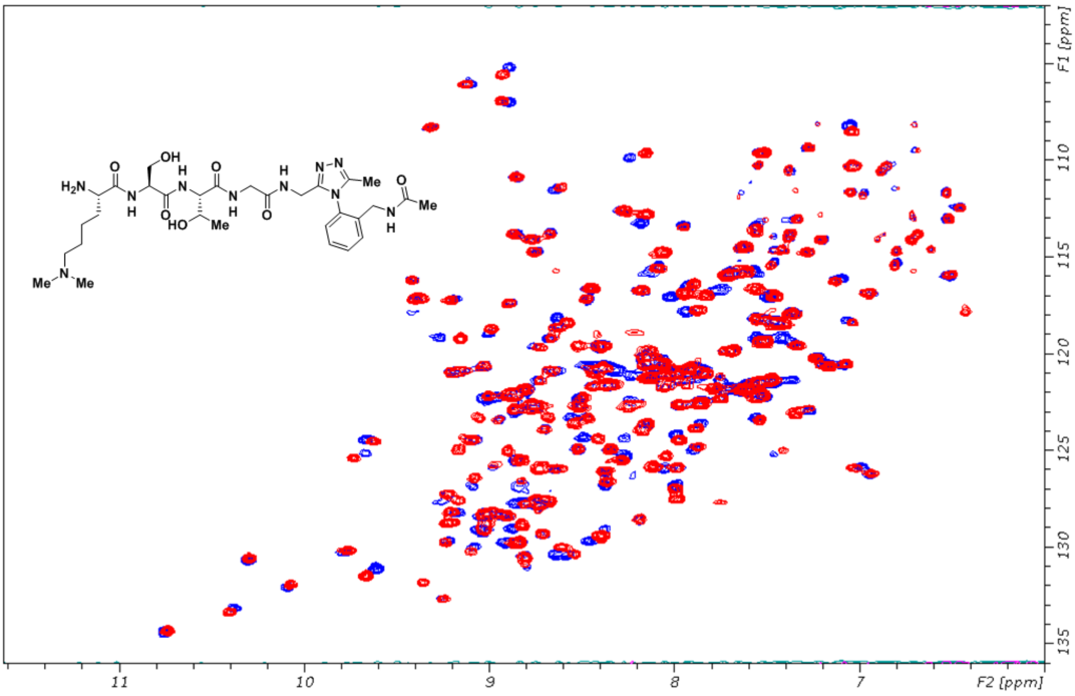

**Supplementary Figure 3**: Commercial Analogues of **1** tested by HSQC NMR

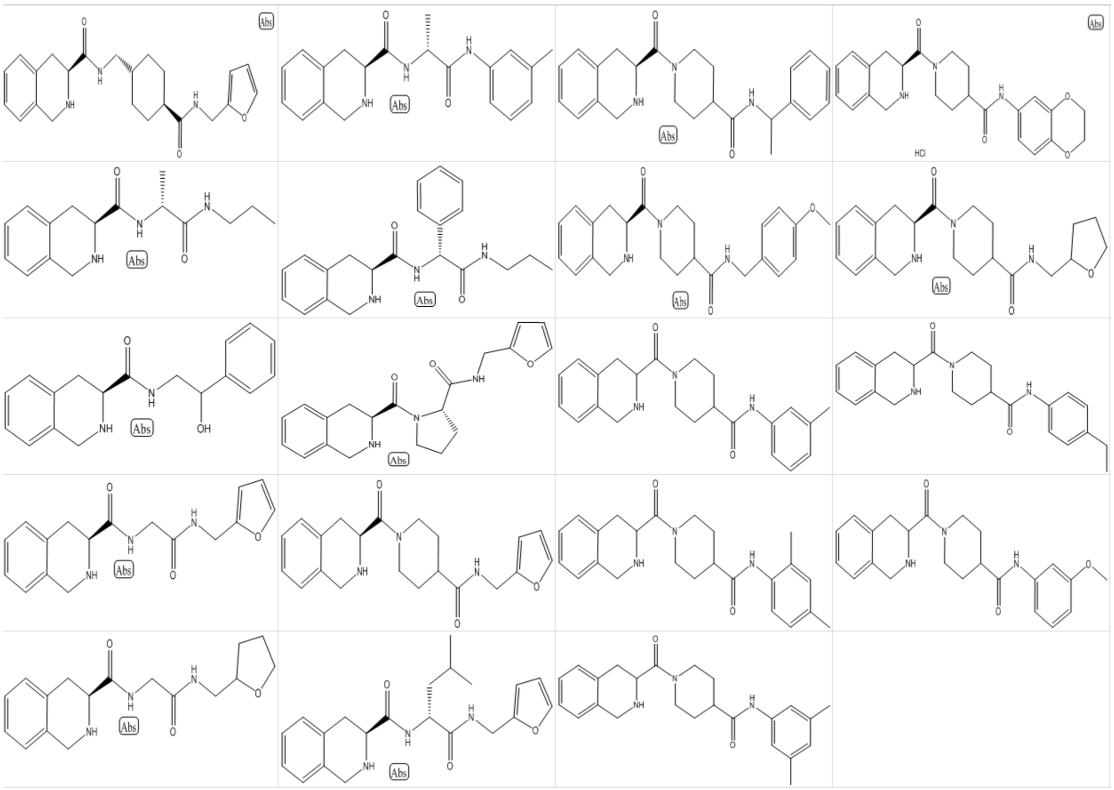

**Supplementary Figure 4**: SPR binding data for compounds **5**, **9a-9d** and **10**.

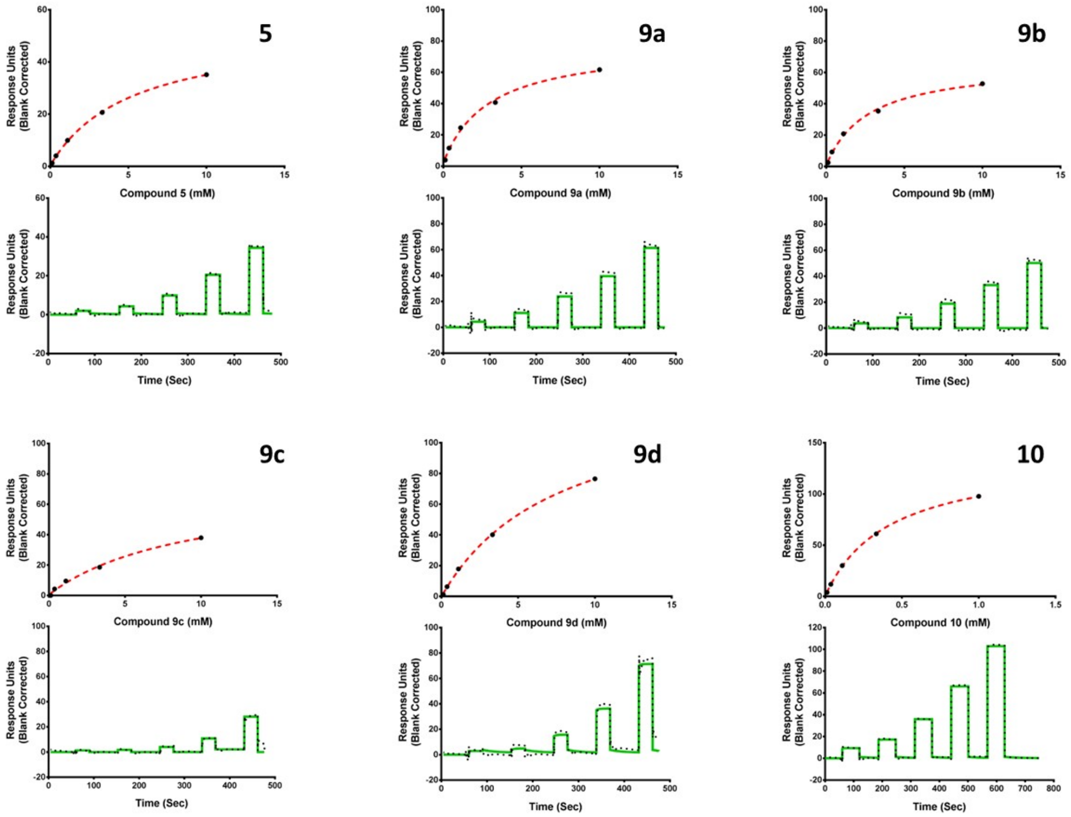


Compounds **5**, **9a, 9b, 9c, 9d** and **10** showed binding affinities with $K_d$ values in a single digit milimolar range. Compound **9c** also showed signs of binding to a control protein which was run in parallel.

**Protein purification procedure**

Purification of SETDB1 Tudor (same procedure for unlabeled and 15N-labeled, different MW)

Purification procedure was based on the Materials and Methods section of http://www.thesgc.org/structures/3DLM

The cell pellets were thawed and resuspended (1 g of cell pellet per 10 mL) in binding buffer (20 mM HEPES pH 7.5, 500 mM NaCl, 5 mM imidazole, 0.5 mM TCEP, 5% glycerol) with protease inhibitor (0.1 mM phenylmethyl sulfonyl fluoride, PMSF). The cell suspension was supplemented with 0.5% (w/v) CHAPS, 5 µl of benzonase (EMD Millipore, cat. no. 70746), protease inhibitor cocktail (Roche) and the cells were sonicated on ice for 5 min total (10 s pulses with 5s interruptions). The lysate was clarified by centrifugation at 20,000 × g, 4 °C, 60 min and the resulting supernatant was filtered through 0.45 µm filter and applied onto 5 mL HisTrap HP column (GE). The column was washed with 10 CV of wash buffer (20 mM HEPES, pH 7.5, 500 mM NaCl, 40 mM imidazole, 0.5 mM TCEP, and 5% glycerol) and the protein was eluted using elution buffer (20 mM HEPES, pH 7.5, 500 mM NaCl, 250 mM imidazole, 0.5 mM TCEP, and 5% glycerol). Next the protein was applied on 26/60 Superdex 200 (GE) column equilibrated with 20 mM HEPES, pH 7.5, 150 mM NaCl, 0.5 mM TCEP at flow rate 2 ml/min. As the final purification step, the protein was purified to homogeneity by ion-exchange chromatography on Source 30S column (10x10) (GE), equilibrated with 20 mM HEPES, pH 7.4, and eluted with linear gradient of NaCl up to 500 mM concentration (20CV). Purification yield was 1 mg of the protein per 1L of culture. The MW (26293.2 Da for unlabeled and 26 616.2 Da for the 15N labeled) of the purified construct was confirmed by LC/MSD TOF (Agilent).

**Purification of BIOH-SETDB1 Expression (used for SPR experiments)**

1. Preparation of Solutions

IMAC lysis buffer: 50mM Tris-HCl, 300 mM NaCl, 5% glycerol, 10 mM imidazole, 0.5 mM TCEP, 0.05% Triton X100 pH 8.0

IMAC wash 1 buffer: 50mM Tris-HCl, 300 mM NaCl, 5% glycerol, 10 mM imidazole, 0.5 mM TCEP, pH 8.0

IMAC wash 2 buffer: 50mM Tris-HCl, 300 mM NaCl, 5% glycerol, 20 mM imidazole, 0.5 mM TCEP, pH 8.0

IMAC elution buffer: 50mM Tris-HCl, 300 mM NaCl, 5% glycerol, 300 mM imidazole, 0.5 mM TCEP, pH 7.5

Gel filtration buffer: 20 mM HEPES, 300 mM NaCl, 5% glycerol, 0.5 mM TCEP, pH 7.5

2. Samples preparation

1). The frozen cell pellets are briefly thawed in warm water. Resuspend the cell pellets by adding Lysis buffer to final volume 100 mL/(per liter culture), Freshly add PMSF and Benzamidin to a working concentration of 1mM and add Benzonase to 5U/ml.

2). Sonication with program nr 1 (5s/7s, 10 min, ~100W) for lysate from 2 L cultivation.

3). The sonicated lysates are centrifuged for 60 min at 16000 rpm in the JLA 16.250 rotor. The soluble fraction is decanted and filtered through 0.45□m filters.

3. Purification

The 5 mL chelating columns and gel filtration columns were equilibrated with IMAC wash 1 buffer and Gel filtration buffer, respectively.

Purification programs:

Chelating 5ml GFS75 2nd wash 20CV level and slope

Chelating 5ml GFS200 2nd wash 20CV level and slope

1. Filtered samples are loaded to the IMAC columns on position 1-4 from sample inlets S1-S4

2. IMAC columns are washed by 10 CV of IMAC wash 1 buffer and 10 CV of IMAC wash 2 buffer.

3. IMAC columns are eluted by 5 CV IMAC elution buffer, and major peak is loaded to gel filtration columns on position 5.

4. The fractions from gel filtration column are checked by SDS-PAGE and pool the pure fractions together. The antigens are further validated by Mass spectrum.

5. The pure antigen samples are diluted or concentrated to 1 mg/ml and make 100 uL aliquots. The aliquots are flash freezed by liquid nitrogen and ship to binders lab with dry ice.

### HPLC chromatograms of final products

RP-HPLC/MS: Low-resolution mass spectra (ESI) were collected on an Agilent Technologies 1200 series HPLC paired to a 6130 Mass Spectrometer. Compounds were resolved on Phenomenex’s Kinetex 2.6u C18 50x4.6mm column at room temperature with a flow of 1 mL/min. The gradient consisted of eluents A (0.1% formic acid in double distilled water) and B (0.1% formic acid in HPLC-grade acetonitrile). ***Method A:*** A linear gradient starting from 5% of B to 95% over 15 min at a flow rate of 1.0 mL/min. ***Method B:*** A linear gradient starting from 5% of B to 95% over 4.00 min at a flow rate of 1.0 mL/min. Stays constant at 95% for 1 min and then returns to 5% over 0.5 min. ***Method C:*** Stays constant at 5% of B for 1.00 min at a flow rate of 1.0 mL/min, followed by a linear gradient to 95% over 8.0 min. Stays constant at 95% of B for 1.00 min and then returns to 5% B over 1.00 min. *Compound 2 (Method A); Compounds 4, 9c, 9d (Method B); Compounds 6-8, 9a, 9b, 9e (Method C).*

### Compounds

**2**

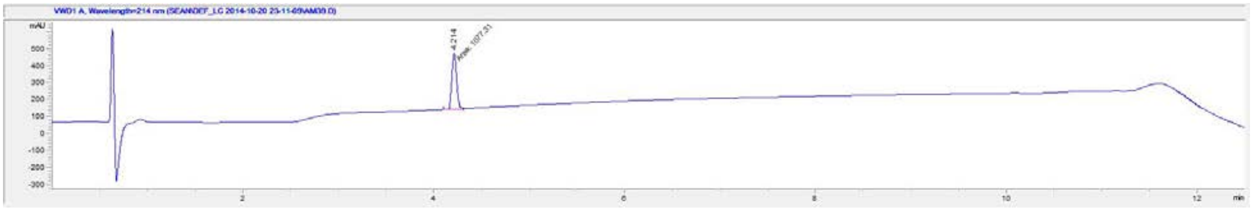

| # | Time | Area | Height | Width | Area% | Symmetry |
|---|---|---|---|---|---|---|
| 1 | 4.214 | 1077.3 | 328.9 | 0.0546 | 100.000 | 0.861 |

**3 : Commercial**

**4**

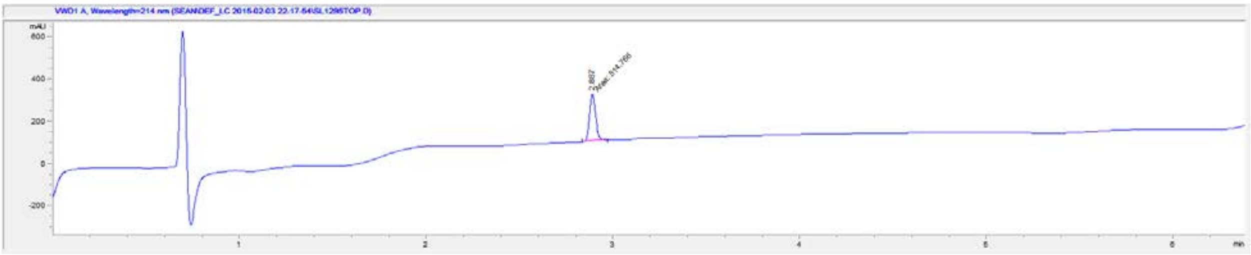

| # | Time | Area | Height | Width | Area% | Symmetry |
|---|---|---|---|---|---|---|
| 1 | 2.887 | 514.8 | 224.6 | 0.0382 | 100.000 | 0.72 |

**5**

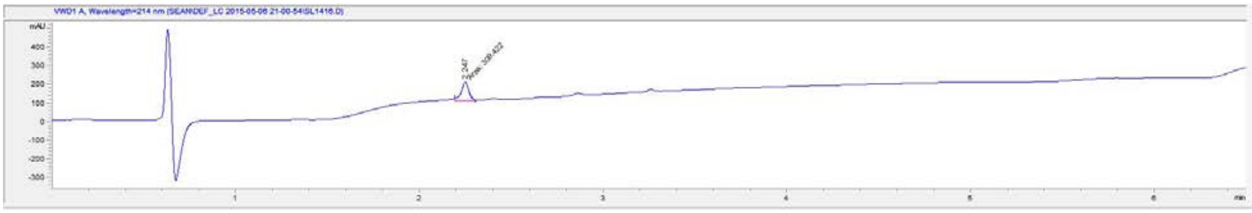

| # | Time | Area | Height | Width | Area% | Symmetry |
|---|---|---|---|---|---|---|
| 1 | 2.247 | 319.5 | 101.7 | 0.0524 | 100.000 | 1.259 |

**6**

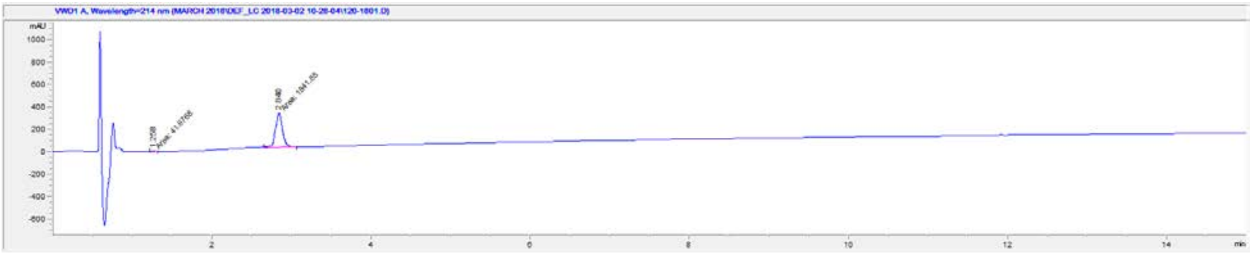

| # | Time | Area | Height | Width | Area% | Symmetry |
|---|---|---|---|---|---|---|
| 1 | 1.258 | 42 | 9.5 | 0.0725 | 2.228 | 0.793 |
| 2 | 2.84 | 1841.8 | 306.6 | 0.1001 | 97.772 | 0.936 |

**7**

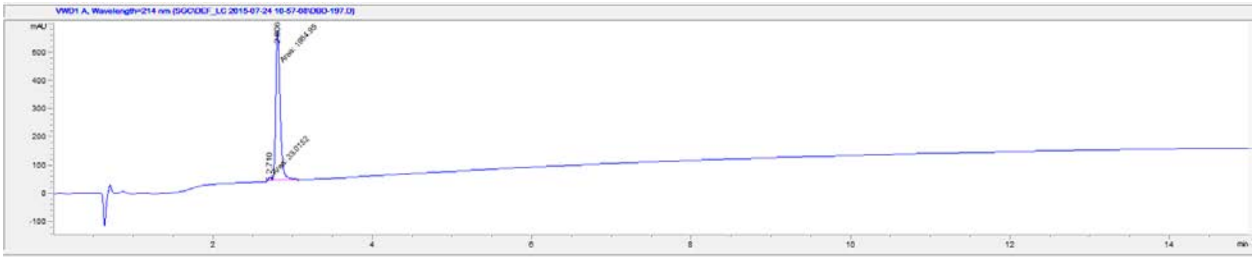

| # | Time | Area | Height | Width | Area% | Symmetry |
|---|---|---|---|---|---|---|
| 1 | 2.71 | 33 | 14.6 | 0.0378 | 1.636 | 0.978 |
| 2 | 2.806 | 1984.9 | 525.9 | 0.0629 | 98.364 | 0.605 |

**8**

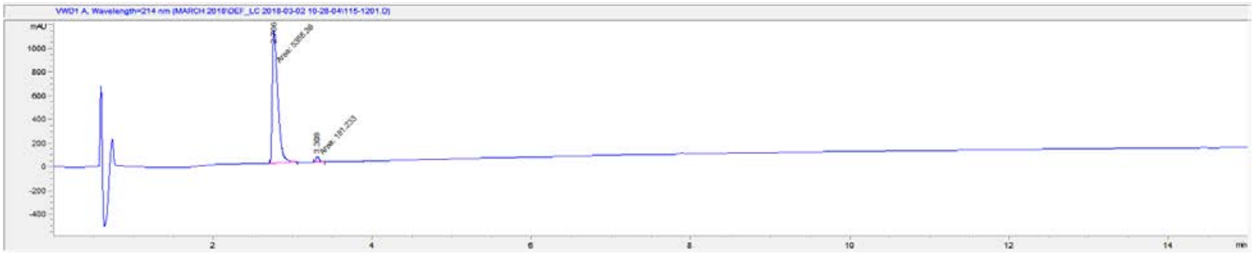


| # | Time | Area | Height | Width | Area% | Symmetry |
|---|---|---|---|---|---|---|
| 1 | 2.766 | 5388.4 | 1120.7 | 0.0801 | 96.746 | 0.389 |
| 2 | 3.309 | 181.2 | 52.6 | 0.0574 | 3.254 | 0.749 |

**9a**

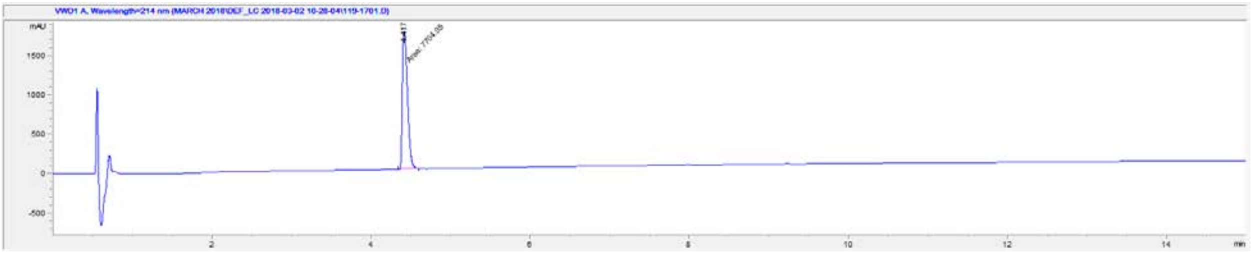


| # | Time | Area | Height | Width | Area% | Symmetry |
|---|---|---|---|---|---|---|
| 1 | 4.417 | 7704.1 | 1743 | 0.0737 | 100.000 | 0.452 |

**9b**

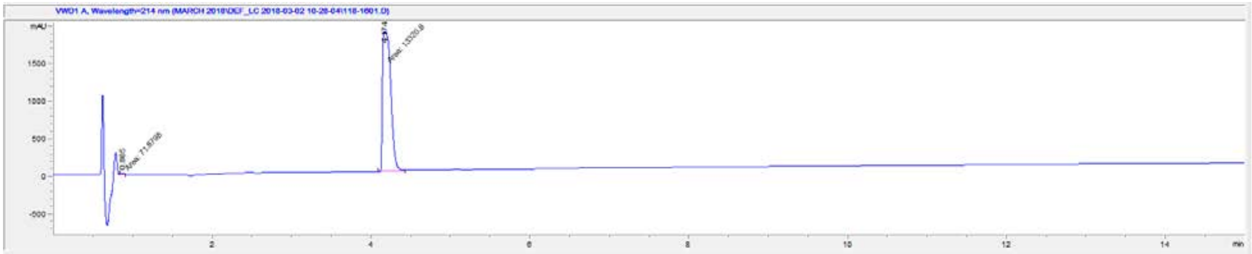

| # | Time | Area | Height | Width | Area% | Symmetry |
|---|---|---|---|---|---|---|
| 1 | 0.865 | 71.9 | 26.8 | 0.0447 | 0.537 | 0.63 |
| 2 | 4.174 | 13320.6 | 1856.1 | 0.1196 | 99.463 | 0.321 |

**9c**

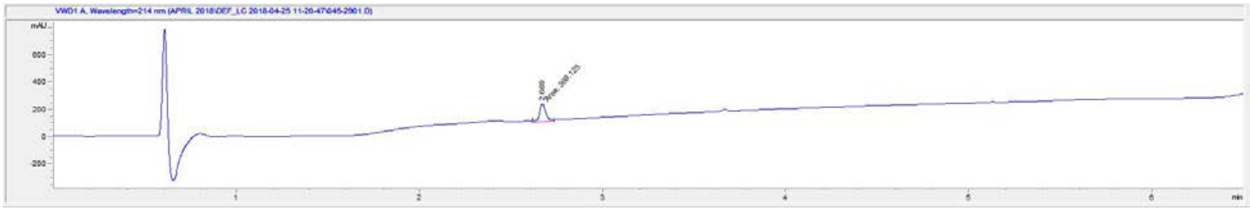

| # | Time | Area | Height | Width | Area% | Symmetry |
|---|---|---|---|---|---|---|
| 1 | 2.669 | 369.1 | 139.4 | 0.0441 | 100.000 | 0.784 |

**9d**

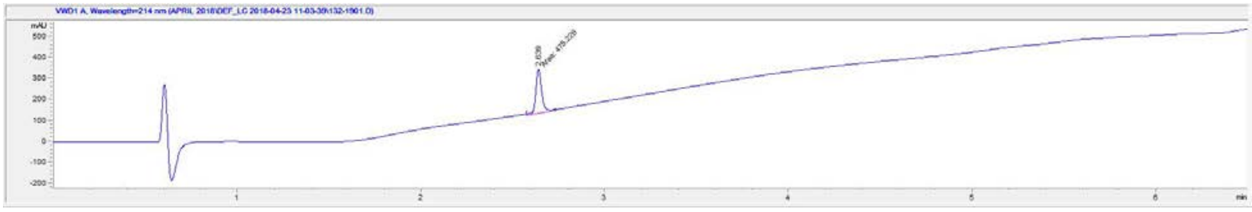

| # | Time | Area | Height | Width | Area% | Symmetry |
|---|---|---|---|---|---|---|
| 1 | 2.639 | 475.2 | 212.9 | 0.0372 | 100.000 | 0.86 |

**9e**

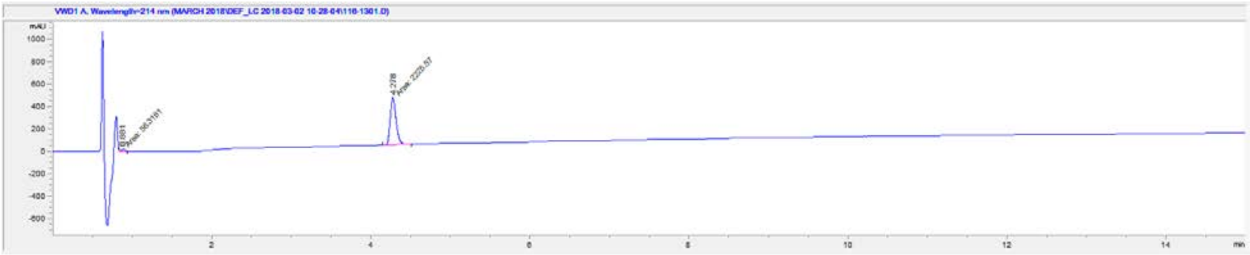

| # | Time | Area | Height | Width | Area% | Symmetry |
|---|---|---|---|---|---|---|
| 1 | 0.881 | 56.3 | 18.8 | 0.0499 | 2.468 | 1.036 |
| 2 | 4.278 | 2225.6 | 425.8 | 0.0871 | 97.532 | 0.732 |

**10**

**98% Pure by 1H NMR spectroscopy. Compound 10 came out in the solvent front of the LC-MS, therefore we used proton NMR to calculate purity**

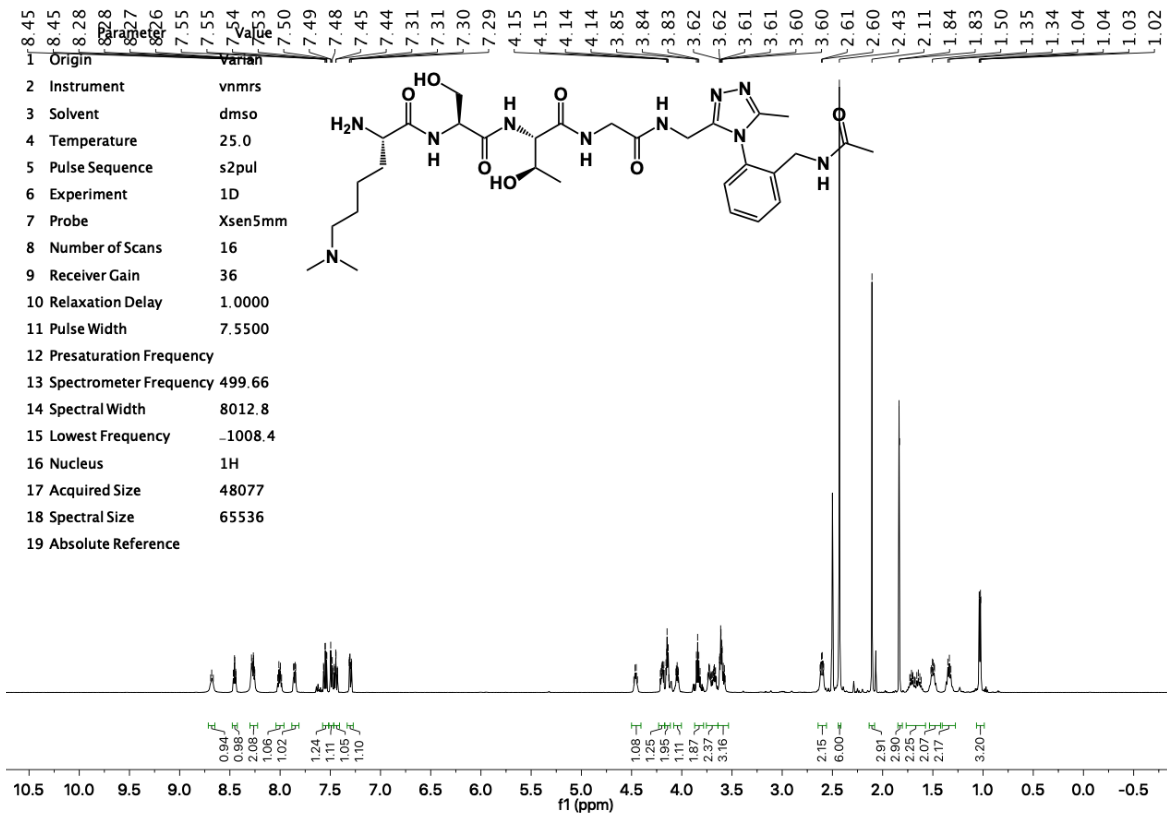

**Compound 10: 13C NMR**

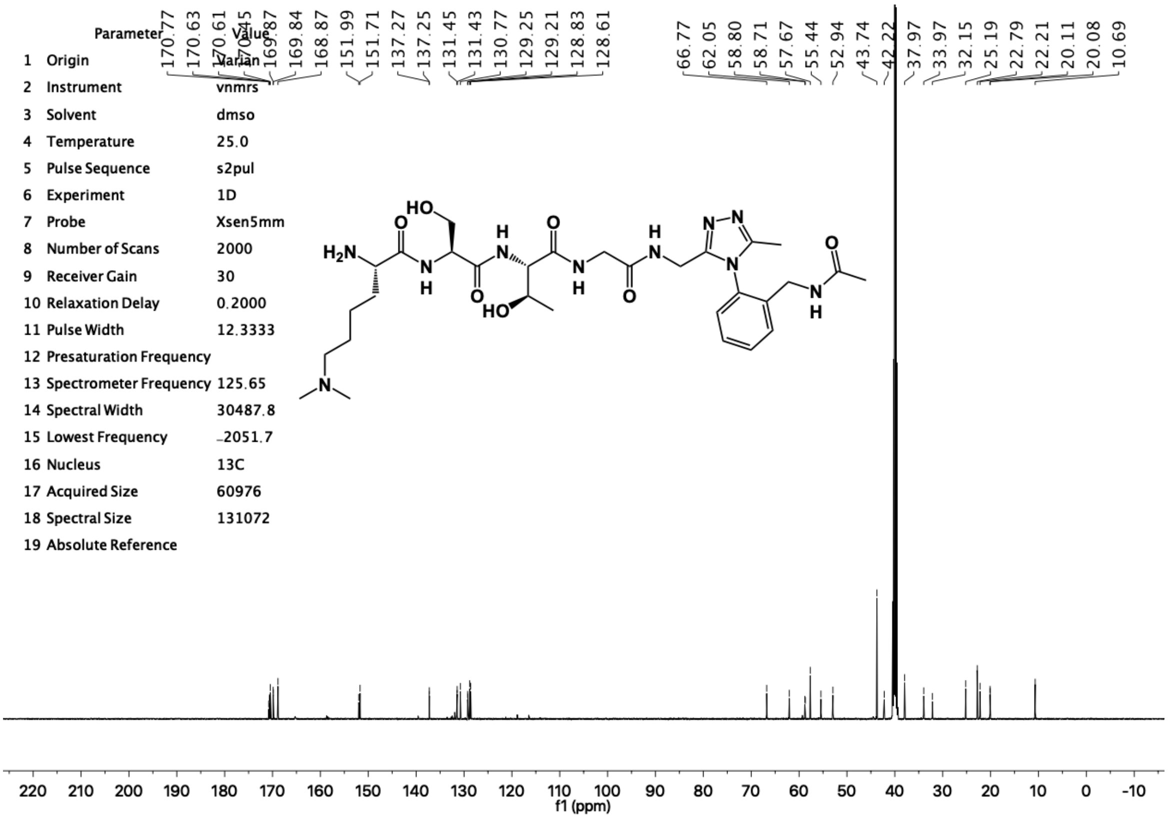
Parameter Value
1 Origin Varian
2 Instrument vnmrs
3 Solvent dmso
4 Temperature 25.0
5 Pulse Sequence s2pul
6 Experiment 1D
7 Probe Xsen5mm
8 Number of Scans 2000
9 Receiver Gain 30
10 Relaxation Delay 0.2000
11 Pulse Width 12.3333
12 Presaturation Frequency
13 Spectrometer Frequency 125.65
14 Spectral Width 30487.8
15 Lowest Frequency -2051.7
16 Nucleus 13C
17 Acquired Size 60976
18 Spectral Size 131072
19 Absolute Reference
170.77
170.63
170.61
170.45
169.87
169.84
168.87
151.99
151.71
137.27
137.25
131.45
131.43
130.77
129.25
129.21
128.83
128.61
66.77
62.05
58.80
58.71
57.67
55.44
52.94
43.74
42.22
37.97
33.97
32.15
25.19
22.79
22.21
20.11
20.08
10.69
f1 (ppm)
